# Sleep promotes aversive value coding in *Drosophila*

**DOI:** 10.1101/2025.07.04.663253

**Authors:** Leonie Kirszenblat, Makoto Someya, Hokto Kazama

**Affiliations:** RIKEN Center for Brain Science, Wako, Saitama, Japan; Graduate School of Arts and Sciences, The University of Tokyo, Meguro-ku, Tokyo, Japan

## Abstract

Sleep is a state of reduced behavioural responsiveness to the external world. Lowered arousal is thought to arise from sensory gating in the brain, yet there is compelling evidence that stimuli can still be processed to some degree. Which aspects of sensory information are processed during sleep remains largely unknown. Here, we perform comprehensive recordings of activity from multiple neuron types during sleep and wakefulness in the *Drosophila* olfactory association center. We find that baseline activity of dopamine neurons (DANs) is distinct between active, rest and sleep states, enabling a decoder to predict when the fly is asleep. Optogenetic inhibition of DANs acutely induces sleep and lowers odor-evoked arousal, suggesting causality between DAN activity and behavior. We then examine odor processing during sleep, and find that whereas coding of odor *identity* in Kenyon cells (KCs) remains intact, coding of odor *value* in DANs is altered. Specifically, DANs encoding positive (attractive) and negative (aversive) values change their odor responses in an opposing manner, producing more aversive odor representations during sleep. Consistent with these changes in neural activity, sleeping flies awoken by odors show reduced approach and increased avoidance behaviour. Taken together, our data reveal that the brain does not simply suppress the propagation of sensory signals during sleep; rather, it utilizes different cell types to retain or modify specific aspects of sensory information. The aversive odor evaluation in DANs may have ethological benefits as it suppresses the motivation to pursue attractive odors during sleep and primes defensive action when aroused by the scent of potential dangers.

## Main

When we sleep, we lose conscious perception of the outside world and our ability to respond to it. This loss of behavioural responsiveness is a defining feature of sleep in all animals^7^. In mammals, reduced arousal is hypothesized to arise from sensory gating in the thalamus that blocks information flow from the thalamus to the cortex^8^. Although a number of studies support this theory, the evidence remains inconclusive^9,10^. It is also clear that sleep does not fully suppress incoming information, as animals can awaken to important cues. For example, humans more likely awaken to their own infant’s cry^11^ and other emotionally or self-relevant stimuli^12^, and mice are aroused more to predator than nonpredatory odors during sleep^4^. Animals can also be aroused by attractive cues, especially if they are unsatiated: a recent study in flies showed that starved individuals awaken more to food-related odors^5^. Apart from arousing an animal, stimulus exposure during sleep can influence learning and memory - a further clue that sensory information must be being processed to some degree^1–3^. Nevertheless, there is still little known about *where*, *how* and *which* aspects of information is processed differently in the brain during sleep^9,10,13–15^. Addressing these questions requires comprehensive monitoring of multiple neuron types across sensory encoding brain areas - a task that has been difficult to achieve in mammals due to its sheer size and complexity.

Here we used the fruit fly, *Drosophila melanogaster*, in which it is possible to image entire brain regions at single cell or cell-type specificity, to directly examine sensory computations during sleep. We focused on the modality of olfaction because the underlying neural circuits and computations have been well described. The mushroom body (MB) is a higher order region for olfactory processing, comprising distinct types of neurons that encode odor identity, assign odor value, and modulate value-driven behaviour^16–20^. Several of these neurons also regulate sleep and arousal^5,20–25^; yet whether their ongoing activity encodes the sleep state, or whether they compute sensory information differently during sleep, has never been observed. In this study, we recorded baseline and odor-evoked activity of multiple cell types throughout the MB during spontaneous sleep. We found that while odor identity representation remains intact, coding of odor value is altered during sleep via coordinated changes in the activity of specific types of DANs. Behavioural responses upon arousal from sleep reflected the changes in neural activity. This reveals cellular- and system-level processing of specific aspects of sensory information during sleep.

## Results

### Baseline activity of MB Kenyon cells and DANs can predict the sleep state

Sleep in *Drosophila* is traditionally defined as a state of immobility lasting 5 min or more, accompanied by reduced arousal to stimuli^26,27^. In mammals, neural correlates of sleep are also used to determine when an animal is sleeping. In *Drosophila*, although rest and sleep have been found to coincide with reduced neural and glial activity^13–15,28^, the identity and activity of neuron types reflecting spontaneous sleep remain unknown.

Therefore, we first sought to identify the neuronal correlates of sleep in *Drosophila*, by performing simultaneous recordings of neural activity and behaviour. A fly walked on an air supported ball under a two-photon microscope, while being tracked with a camera and software to infer walking manoeuvres from the movement of the ball^29^ (Fig. 1a). We found that the distribution of estimated walking speeds was distinct between manually annotated active and inactive states, such that a threshold value of 1.2 mm/s could be used to automatically classify the behavioural state (Fig. 1b). Using this threshold, we determined periods of activity, rest (inactivity < 1 min), sleep (inactivity > 5 min) and intermediate periods of inactivity (Fig. 1c). Flies were often aroused to the onset of the laser scanning when they were in the active state, but aroused much less frequently after 5 min of inactivity - fulfilling the criteria for sleep in *Drosophila* in both freely-walking^26,27^ as well as tethered preparations^14^ (Fig. 1d). Notably, flies showed significantly reduced arousal even from earlier timepoints (1-5 min inactive), suggesting that they may sometimes enter a sleep-like state more rapidly^15,30^. In contrast, flies that had been inactive for less than 1 min were significantly more arousable (Fig. 1d), consistent with a report that only after 1 min do they display pre-sleep behaviour ^31^.

**Figure 1.**
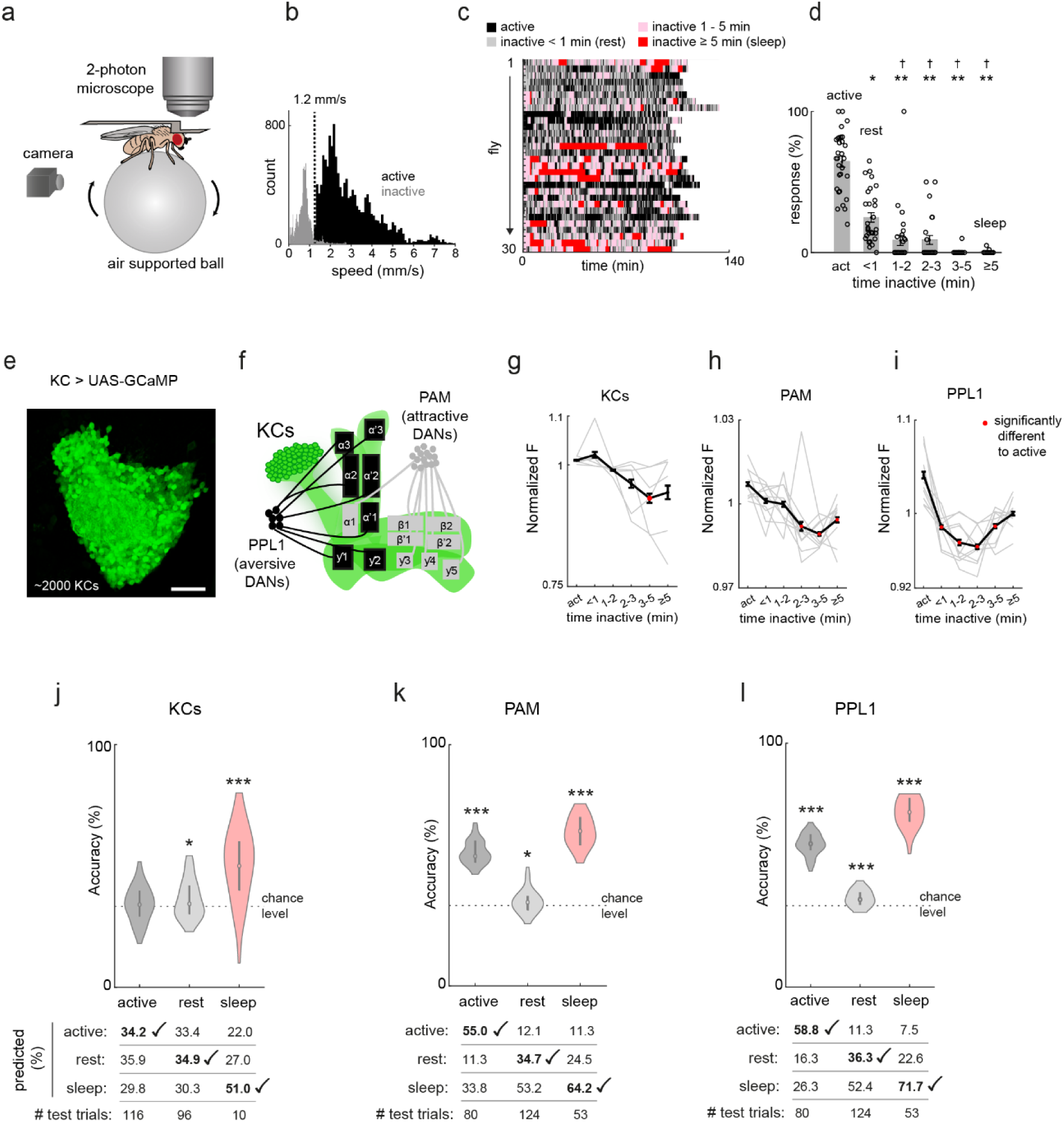
DAN and KC baseline activity can predict behavioural states. (**a**) Setup for simultaneous recording of behaviour and neural activity. (**b**) Histogram of walking speeds during annotated behavioural periods, indicating when the fly was active (black) or inactive (grey) (n = 4 flies). (**c**) Raster plot showing transitions between behavioural states across individual experiments (each row corresponds to one fly). Black indicates active movement, light grey indicates short rest (inactivity < 1 min), red indicates sleep (inactivity ≥ 5 min), and pale pink represents intermediate inactivity (1–5 min of inactivity). (**d**) Flies are less easily aroused by the start of the scanning laser after prolonged periods of inactivity. Responses to the scan start are shown for each fly. When flies were inactive prior to the stimulus, they showed reduced responses, and a further reduction after 1 min of inactivity (*p < 0.05, **p < 1e-07, compared to the active state, †p < 0.05 compared to rest, Kruskal-Wallis test with Dunn-Sidak correction for multiple comparisons, n = 30 flies). (**e**) Volumetric image of ∼2000 KC somata visualized with the Ca^2+^ indicator UAS-GCaMP6f. Scale bar represents 25μm. (**f**) Schema showing regions imaged in the MB. KC somata and DANs conveying positive sensory signals (PAMs) and negative sensory signals (PPL1s) were imaged in each of the 15 MB compartments. (**g-i**) Average baseline activity of KCs (**g**), PAMs (**h**) and PPL1s (**i**) recorded during periods of activity (“act”) or after increasing periods of inactivity. Each line represents data from a single fly across the three behavioural states, with black line and error bar representing mean and s.e.m. Means are highlighted in red when the value was significantly different from the active states (Kruskal-Wallis test, p < 0.05, after Dunn-Sidak correction for multiple comparisons). n = 6 flies for KCs and 10 flies for PAM/PPL1s. (**j-l**) KC and DAN (PAM or PPL1) baseline activity can predict sleep above chance level, although the accuracy is higher for PAMs and PPL1s compared to KCs. Each distribution represents the accuracy of prediction (percentage correct) across 50 decoding models trained and tested with different sets of trials (see Methods). Table below graph shows the number of test trials and the predicted results (*p < 0.017, ***p < 1e−08, one-tailed t-test with Bonferroni correction compared to chance level of 100/3%). n = 10 flies for DAN analysis and 6 flies for KC analysis.

We first recorded from all ∼2,000 somata of Kenyon cells (KCs) (Fig. 1e), the principal intrinsic neurons of the MB^16,32^ (Fig. 1f). Baseline activity was imaged for ∼6 s every minute in each experiment, which typically lasted for ∼100 min. Ca^2+^ levels averaged across KCs did not change in the first few minutes of inactivity but gradually declined thereafter, similar to a previous report^13^ (Fig. 1g).

We next recorded from all the DANs in the MB. Each type of DAN innervates one or a few of the 15 MB compartments^33^ and belongs to either PAM or PPL1 clusters, which convey information about positive or negative sensory signals, respectively^34–36^ (Fig. 1f). GCaMP signals in individual MB compartments were extracted by registering a three-dimensional template of the compartments to individual imaged brains^17,33^ (Supplementary Fig. 1). Similar to KCs, baseline activity of PAM neurons declined the longer the fly had been resting (Fig. 1h), while that of PPL1 neurons dropped more sharply even within a minute of resting, with a slight rise again after 3 min (Fig. 1i). This suggests that DANs show distinct dynamics that reflect behavioural states of activity, rest and sleep.

These state-dependent dynamics suggest that KC and especially DAN baseline activity may suffice to predict when flies are in any of the above behavioural states. To examine this, we trained a decoder to classify the states into active, rest and sleep using the baseline activity of KC, PAM or PPL1 in a subset of trials, and tested whether it could predict the state on withheld trials. We found that sleep could be identified above chance level (33%) with a mean accuracy of 51% for KCs (Fig. 1j). Prediction of the sleep state was even more accurate for DANs, with a mean accuracy of 64% and 72% for PAMs and PPL1s (Fig. 1k,l). The active state was also identified significantly above chance level for PAM and PPL1s, but not for KCs. In contrast, the rest state was more difficult to classify, often being misclassified as sleep (Fig. 1k,l). Thus, DAN activity during sleep sometimes resembles that during rest, consistent with reports that flies have different sleep stages that comprise ‘quiet’ and ‘active’ sleep^15,30,37–39^. Together, our results show that DAN and, to a lesser degree, KC activity levels, reflect whether flies are in the sleep state, which is distinct from rest.

### Optogenetic inhibition of DANs induces sleep

We next asked whether DAN activity causally regulates arousal levels and sleep. Previous studies have shown that mutations in the dopamine pathway modulate sleep in *Drosophila*^40–44^. Temperature-induced activation of subsets of MB DANs mostly promote wakefulness^25^. On the other hand, inhibition of DANs increases sleep^25,43,45^, although the use of chronic inhibition may have induced developmental side effects. Furthermore, the effect of DAN inhibition on arousal to stimuli remains unknown. Since we observed that both PAM and PPL1 DANs lower their baseline activity during and preceding sleep (Fig. 1h,i), we hypothesized that suppressing activity in both types of DANs would promote sleep and reduce stimulus-evoked arousal. In addition, as the decline in PAM baseline activity continued during sleep (Fig. 1h), inhibition of PAMs was expected to exert a stronger effect.

To test these hypotheses, we optogenetically inhibited DANs by expressing a light-gated ion channel, GtACR1^46^. This method allowed us to precisely control the timing and intensity of inhibition, without the developmental effects of chronic manipulation or the physiological effects of heat. We examined odor-evoked behaviour while applying two different intensities of light to activate GtACR expressed with either *TH-Gal4* or *DDC-Gal4* that label largely non-overlapping sets of DANs^35^ (Fig. 2a); *TH-Gal4* labels PPL1s with minimal PAMs whereas *DDC-Gal4* heavily labels PAMs^34,35^. We found that inhibition of both types of DANs promotes sleep with a more salient effect in PAMs, especially at the stronger LED intensity (Fig. 2a,b). This is consistent with the result that baseline activity was suppressed in both types of DANs during rest and sleep (Fig. 1h,i). The weaker effect of PPL1 manipulation may reflect the rise in its activity during sleep, following the decline during the first several minutes of behavioral inactivity (Fig. 1i).

**Figure 2.**
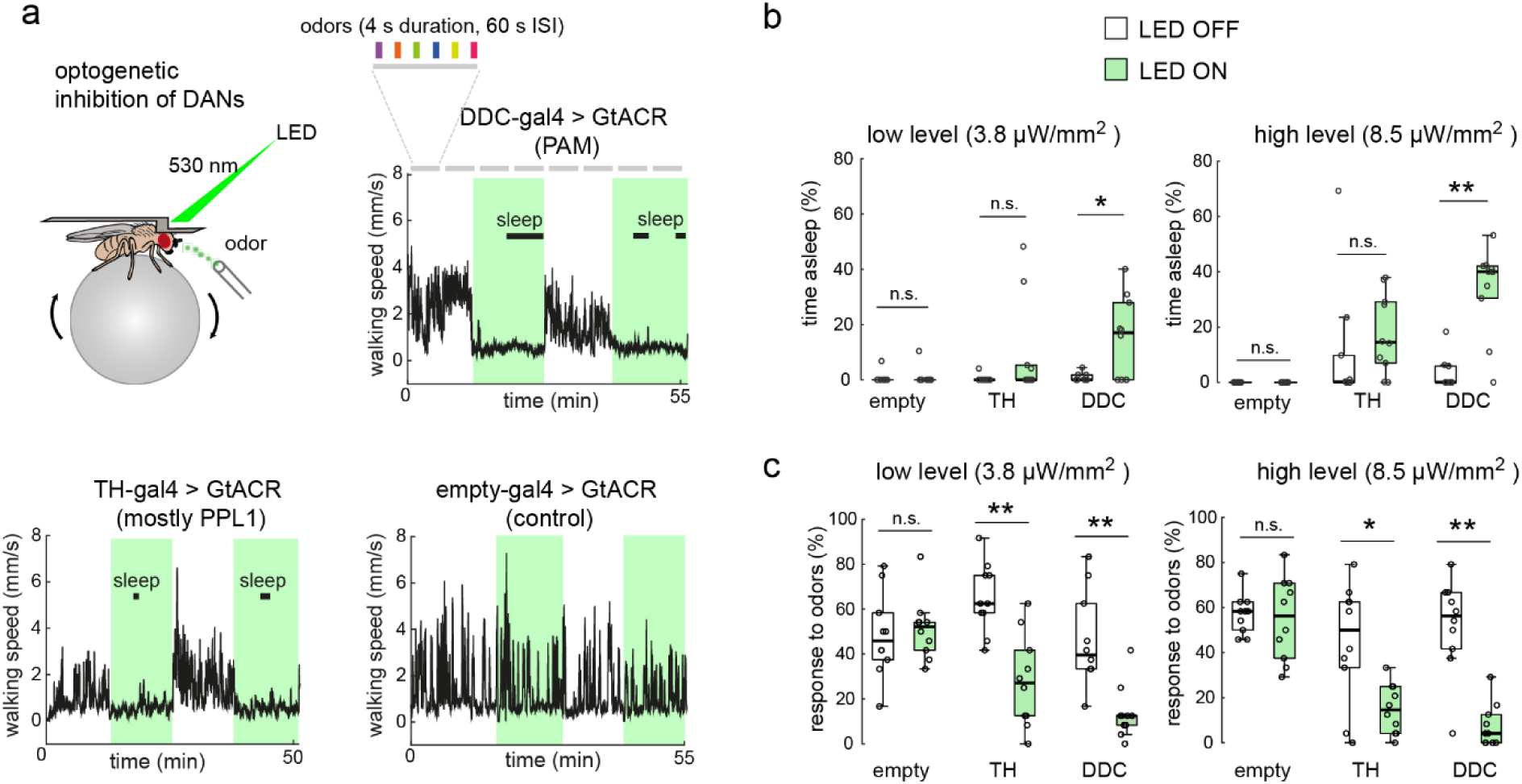
Optogenetic inhibition of dopamine neurons promotes sleep and reduces odor-evoked arousal. (**a**) DANs were optogenetically inhibited by expressing GtACR using *TH-Gal4* (labelling PPL1s with minimal PAMs) or *DDC-Gal4* (heavily labelling PAMs). Sleep and arousal to 6 different odors were measured across two sets of contiguous control blocks (LED OFF) and two sets of inhibition blocks (LED ON). Walking speed and sleep epochs during LED on and off periods are shown for representative individuals with different genotypes. (**b**) Percentage time spent asleep during LED OFF and ON periods for two different light intensities (each dot represents a fly). (**c**) Percentage of trials in which flies responded to odors during LED OFF and ON periods. Boxplots show the median (central line), the interquartile range (box edges), and whiskers (the most extreme datapoints within 1.5 times the interquartile range from the lower and upper quartiles), with circles showing the percentage of odor trials to which each fly showed a significant walking response (see Methods). *p < 0.05, **p < 0.01, Wilcoxon signed-rank test, n = 10 flies per genotype).

We further examined if arousal level is also modulated, by presenting flies with different odors. We found that arousal to odors was strongly suppressed by both PAM and PPL1 inhibition (Fig. 2c). Although loss of dopaminergic signalling can lead to deficits in motor control^47,48^, we quantified walking behaviour in trials where flies significantly responded to odors, and found no differences in walking speed during DAN inhibition (Supplementary Fig. 2). Thus, acute inhibition of DAN activity reduces arousal, which is necessary for sleep onset and maintenance.

### Odor identity coding is preserved during sleep

A major task of the olfactory system is to identify and discriminate different odors, yet how this is modulated by arousal state is poorly understood. In *Drosophila*, odor identity is encoded by KCs^16,32^. To investigate whether sleep alters odor identity coding, we recorded responses to 15 odors from all ∼2,000 KC somata with Ca^2+^ imaging^18^ in tethered walking flies (Fig. 3a,b, Supplementary Video1). During imaging, we observed that arousal to odors depended on the behavioural state: flies responded strongly to odors when they were active, slightly less so when they had been inactive for < 1 min (‘rest’), and even less after longer periods of inactivity (Fig. 3c, Supplementary Fig. 3a,b).

**Figure 3.**
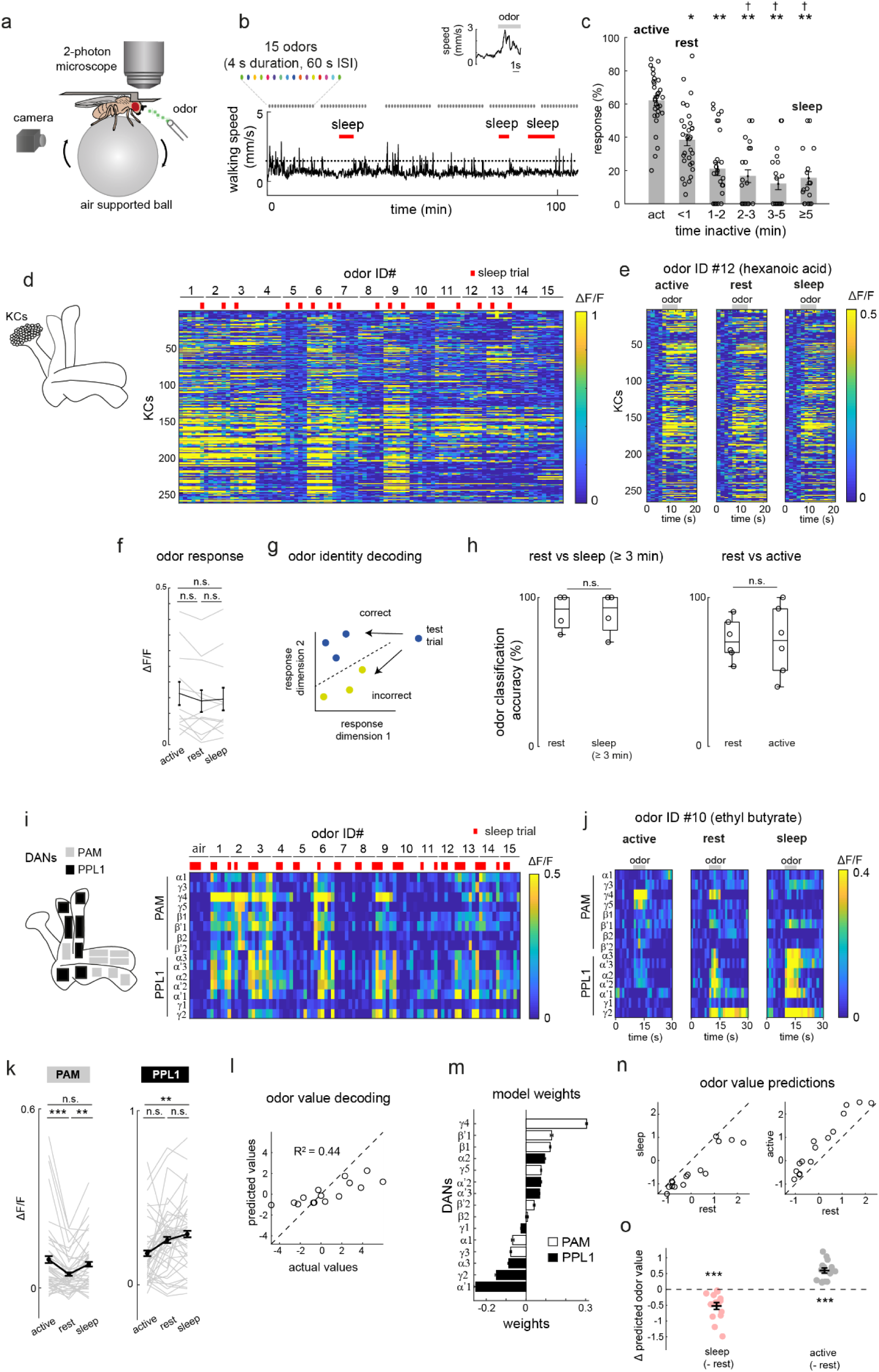
Sleep alters odor value coding in DANs. (**a**) Setup for simultaneous recording of behaviour and brain activity during odor application. (**b**) Walking behaviour during repeated odor stimulation in an example fly. Periods of sleep (inactivity ≥ 5 min) during odor application are marked with the red bar. Colored dots at the top indicate that 15 different odors plus one control stimulus (air) were applied in each block with 4 s duration and 60 s inter-stimulus interval (ISI). Odors are listed in Table S1. The inset graph shows an example of a fly responding to an odor with a significant increase in walking speed. (**c**) Arousal to odors (the percentage of trials in which flies showed a significant walking response, see Methods) depended on time inactive prior to odor application, and was significantly reduced during sleep (inactive for at least 5 min). Asterisks indicate differences from the active state while crosses indicate differences to the ‘rest’ state (when flies have been inactive for less than 1 minute). Mean and s.e.m. are shown, with circles representing each fly (n = 30 flies, *p < 0.05, **p < 1e−07, compared to the active state, † p < 0.05 compared to the rest state, Kruskal-Wallis test followed by Dunn-Sidak correction for multiple comparisons). **(d)** Average odor responses of KCs in an example fly. Red ticks indicate trials in which the fly was asleep. Trials are sorted by the odor identity. KCs that did not respond significantly to any of the tested odors are not shown. Diagram shows the recorded region in the MB (KC somata). (**e**) Time course of example odor responses during active, rest and sleep trials in the same fly. (**f**) Mean odor responses in each state (n = 12 odor matched trials from 5 flies for each state (see Methods), no significant differences between states with Wilcoxon signed-rank test and Bonferroni correction for multiple comparisons: p = 0.18 for active vs sleep, 0.064 for active vs rest and 0.47 for rest vs sleep). Each data point represents a single trial. (**g**) Schematic showing odor identity decoding with linear discriminant analysis (LDA). A model was trained on rest data to separate odors of different identity and the classification accuracy was assessed using the withheld trials from the training state (rest) and test states (sleep or active). The schema illustrates a simple example of discrimination of two odors; in our experiments, up to 15 odors were separated in a high dimensional space. (**h**) Odor identity decoding using KC responses was accurate with no differences between rest vs sleep and rest vs active states (n = 6 trials, 4 flies, p = 1.0 for rest vs sleep, and n = 19 trials, 6 flies, p = 0.93 for rest vs active, Wilcoxon signed-rank test). Boxplots show the median (central line), the interquartile range (box edges), and whiskers (the most extreme datapoints within 1.5 times the interquartile range from the lower and upper quartiles), with circles showing the accuracy level for each fly. (**i**) Average odor responses of DANs in an example fly. Red ticks indicate trials in which the fly was asleep. Trials are sorted by the odor identity. Diagram shows the positions of recorded PAM and PPL1 DANs. (**j**) Time course of example odor responses (to ethyl butyrate) during active, rest and sleep states in the same fly. (**k**) Odor responses averaged across PAM and PPL1 compartments in active, rest and sleep states. Each line represents fly-odor matched data across the three behavioural states, with black line and error bar representing mean and s.e.m. **p < 0.01, ***p < 0.001, Wilcoxon signed-rank tests with Bonferroni correction for multiple comparisons (n = 41 trials, 10 flies). (**l**) A decoding model trained on rest trials predicted odor value from DAN responses. Each dot represents the mean predicted value for each odor, plotted against the actual value. (**m**) Model weights for individual compartments (mean and s.e.m.). (**n**) The model in (**l**) was used to predict odor value in withheld trials where flies were either resting, asleep or active. Here, odor value predictions during rest are plotted against those during sleep or active states. (**o**) Odor values are predicted more negatively during sleep, and more positively during the active state. Each dot shows the difference between mean predicted value during rest and sleep (red dots), or rest and active (grey dots), for each odor (***p < 0.0003, one sample t-test, significantly different to zero).

We found that distinct groups of KCs responded significantly to different odors during periods of activity, rest and sleep (Fig. 3d,e). The average magnitude as well as the pattern of population responses remained relatively stable across these three behavioural states (Fig. 3e, f, Supplementary Fig. 4a,b). To quantify whether odor identity coding was similar between the behavioural states, we employed linear discriminant analysis. In our initial analysis, we defined sleep as when a fly had been inactive for at least 3 min, because this condition coincided with reduced arousal (Fig. 1d, Fig. 3c) and reduced neural activity (Fig. 1h,i see the 3 - 5 min bin) thus resembling sleep. This is consistent with a previous study in freely moving flies, which also proposed that sleep occurs after only 3 min of quiescence^31^. A decoder was trained with rest trials to separate up to 15 odors and tested whether it could correctly identify the odors applied during the withheld rest or sleep trials (Fig. 3g). Surprisingly, odors could be accurately identified even during sleep, with no significant difference to the rest state (Fig. 3h). We also found no difference in decoding accuracy between active and resting states (Fig. 3h). Lastly, a similar result was obtained when we used the more stringent 5-min criterion for sleep; odor identity coding remained accurate during sleep and was no different to wakefulness (active or rest) (Supplementary Fig. 4c). This shows that odor identity coding in KCs is preserved during sleep.

### Odor value coding is altered during sleep

As animals respond more to odors when they find them strongly aversive or attractive, we hypothesized that sleep may suppress odor-evoked behavioural responses by altering the processing of odor *value*. We previously determined the innate value of a large panel of odors by the degree to which they elicited attractive or aversive behaviour^49^, and also showed that these values are represented by DANs innervating the MB^17^.

To assess whether odor value coding is altered during sleep, we recorded odor responses from all the DANs innervating the MB (Fig. 3i, Supplementary Fig. 1). Unlike KCs, DAN odor responses appeared less consistent across trials, hinting that responses may be modulated by sleep (Fig. 3i). For instance, responses to an attractive odor in an example fly show that the major site of activation is shifted from PAM to PPL1 compartments during rest, and this trend becomes even stronger during sleep (Fig. 3j). Indeed, we observed distinct changes in odor responses of PAMs and PPL1s during sleep: whereas average PAM responses were highest when the fly was active and reduced during rest, average PPL1 responses were lowest when the fly was active and increased during rest and even further during sleep (Fig. 3k). A similar pattern of changes could be seen across compartments (Supplementary Fig. 5a,b, Table S2). Furthermore, reduced odor responses during sleep were seen in PAM compartments whose baseline activity during wakefulness encoded forward movement, whereas increased odor responses during sleep were seen in PPL1 compartments encoding backward movement (Supplementary Fig. 6a,b, Table S2). Together, these results suggest that DANs might encode odors more aversively during sleep, which we quantify rigorously below.

As sleep is modulated by GABAergic pathways in the MB^21,23^, we wondered whether these compartment specific changes in neural activity during sleep are coordinated by the GABAergic anterior paired lateral neuron (APL). APL innervates the entire MB (Supplementary Fig. 7a) and is known to provide local or global inhibition^50–52^. However, APL odor responses remained unaffected by behavioural states (Supplementary Fig. 7b,c). Furthermore, APL did not show any increase in baseline activity during rest or sleep (Supplementary Fig. 7d), suggesting that APL has no role in altering odor values in DANs.

To test whether odors become more aversive during sleep, we took a decoding approach and asked whether DAN population activity during active, rest and sleep states differentially predict odor values. We assessed how flies evaluate odors in our tethered walking paradigm by examining the degree to which they walk forward to odors (an attractive response), and defining a value index ranking odors from most attractive to most aversive (Supplementary Fig. 8a). The value index was significantly correlated with the index derived from flying flies^17,49^, suggesting that odor value is similarly evaluated during flight and walking (Supplementary Fig. 8b). We then trained a decoding model on DAN odor responses in rest trials using partial least-squares regression, and tested if it could predict the value of odor presented in withheld rest trials based on DAN responses. The odor value could be accurately predicted (coefficient of determination, R^2^ = 0.44), with PAM and PPL1 DANs typically having the strongest positive and negative weights, as expected (a strong positive weight indicates that the odor response in that compartment contributes more positively to the predicted value, whereas a strong negative weight contributes more negatively) (Fig. 3l,m). Crucially, when we tested with the withheld sleep or active trials, the predicted odor values were mostly negatively shifted for sleep predictions, and positively shifted for active state predictions (Fig. 3n,o). These results show that odors are represented as more attractive during the active behavioural state and as more aversive during sleep.

### Odors are more negatively valued during sleep

We next examined whether this sleep-dependent shift in DAN odor value coding is reflected in the animals’ behaviour. One challenge that arises here is that sleep, by definition, is characterised by a lack of behavioural response. To circumvent this, we took advantage of trials in which flies were awoken by odors. We focused on trials in which flies were stationary prior to odor onset either in the rest or sleep state, and in which they responded to odors with a significant increase in walking speed. This enabled us to examine the nature of the behavioural response (i.e. the degree of approach or avoidance-related behaviour) during sleep or rest. Due to our observation that DANs encoded odors more aversively during sleep, our hypothesis was that behaviour would be biased towards avoidance in sleep as compared to resting trials. As expected, whereas forward velocity towards attractive odors rapidly increased upon odor onset in resting flies, it increased only slowly with a significantly lower mean value in sleeping flies (Fig. 4a,b). Forward velocity in response to neutral and aversive odors was less affected in the mean value (Fig. 4a,b), but it was significantly lower in the maximum value for all odors during sleep, suggesting that flies show reduced instances of approach (Supplementary Fig. 9a). Forward turning was also reduced when awoken from sleep (Supplementary Fig. 9b). Furthermore, aversive odors triggered a more persistent avoidance response in flies awoken from sleep; flies continued to show backward walking (i.e. negative velocities) 2 s after odor onset (Fig. 4b, mean forward velocity = −1.0 ± 0.7 (s.e.m.) mm/s vs 1.1 ± 0.7 mm/s in sleep vs awake, resting flies, p = 0.043, Wilcoxon sign-rank test). The reduced approach and more persistent avoidance upon arousal match the shift in DAN odor value coding during sleep.

**Figure 4.**
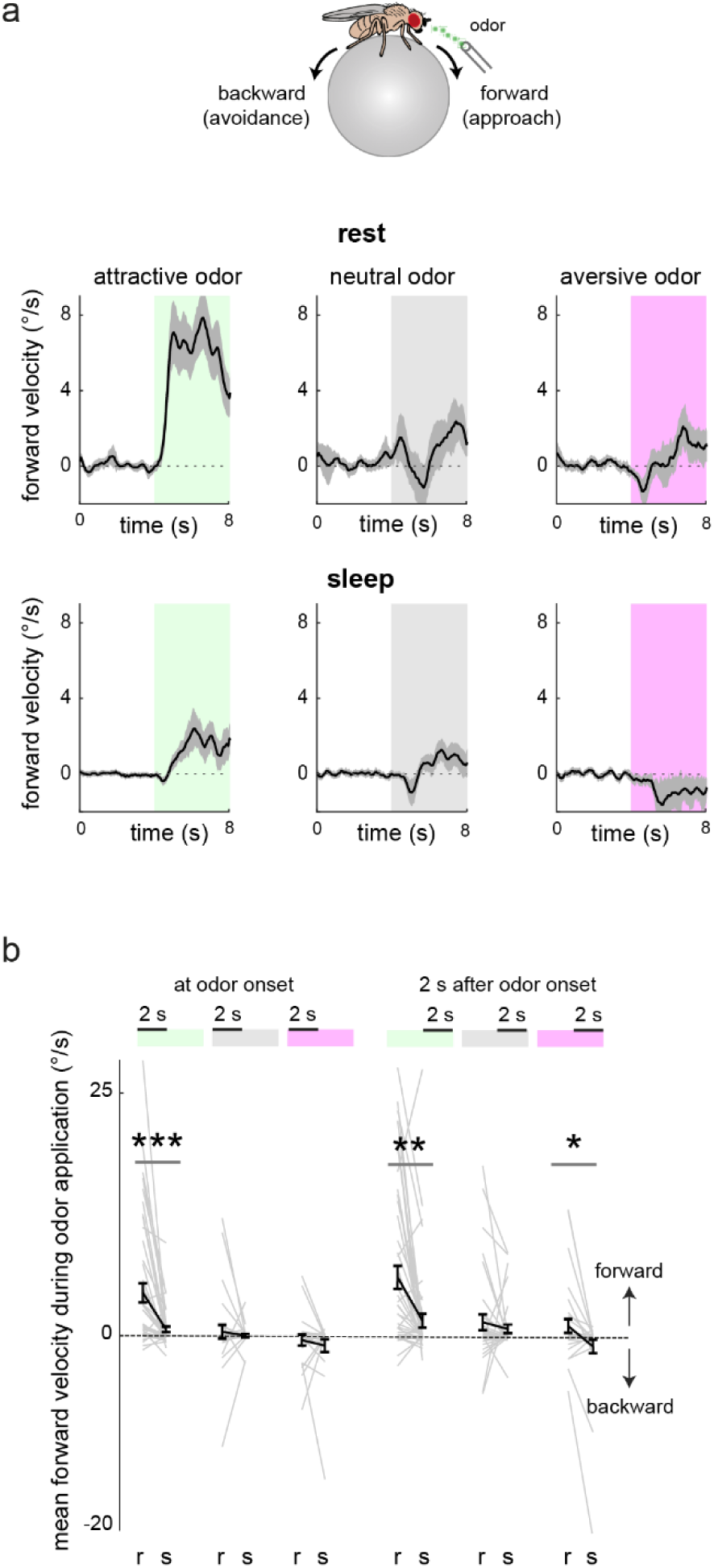
Odors are behaviourally more negatively valued during sleep. (**a**) Mean forward velocity of flies that were awake and inactive, or asleep at the time of odor onset. Only responses with significant changes in velocity were analyzed (i.e. flies were aroused). Mean and s.e.m. are plotted (0.3 s moving average) with odor application period indicated by the shaded region. N= 49, 36 and 32 fly-odor matched trials for attractive, neutral and aversive odor response graphs, respectively. (**b**) Mean forward velocity during odor application for rest (‘r’) and sleep (‘s’) groups in (**a**) either during the first or the next 2 s after odor onset (*p < 0.05, **p < 0.001, ***p < 000001, Wilcoxon signed-rank test).

## Discussion

Our data reveal that sleep does not simply shut out the world using ‘gates’ in the brain. Instead, different aspects of incoming signals are either preserved or altered at distinct nodes in the circuit. We found that whereas odor identity coding in KCs remained intact, odor value coding in DANs was altered such that odors were represented more aversively during sleep. Importantly, the shift in representation was reflected in behavioural responses upon sensory arousal.

Gaining these insights into sensory processing during sleep required overcoming four major technical challenges: recording neural activity 1) at single cell or cell-type resolution, 2) comprehensively from brain regions, 3) through periods of spontaneous sleep and wakefulness, and 4) while applying stimuli without arousing the animal. In mammals, most studies have recorded brain activity during sleep with electroencephalography or intracranial local field potential (LFP) recordings, both of which measure the summed electrical activity of multiple neurons. In cases where sensory-cue evoked unit activity was captured, the exact identity of the neurons was unknown^9^. Although Ca^2+^ and voltage imaging has allowed recording from single and specific types of cells, probing sensory responses during sleep without inducing arousal remained an obstacle. In *Drosophila*, ex vivo recordings have been used to investigate sleep pressure or circadian time, and in vivo recordings have relied on artificially induced sleep^22,53,54^. Neural recordings during spontaneous sleep are scarce - particularly under sensory stimulation - with one study measuring LFP^55^ and another study sampling a limited number of neurons^13^. Here, we overcame these obstacles, investigating the dynamics of multiple cell types throughout the MB, comparing odor responses during sleep and wakefulness. This approach was crucial to go beyond the study of individual neurons to reveal the changes in sensory computations being performed by each neuronal population.

Our results also demonstrate that ongoing DAN activity can predict periods when the fly is asleep, establishing it as a neurophysiological marker of sleep. Rest periods were less accurately predicted, often being misclassified as sleep, suggesting that flies may enter a sleep-like state even after short periods of rest. Although the active state was better classified, it was also occasionally mistaken as sleep. One possibility is that these periods of DAN activity resembling the sleep state are akin to ‘local sleep’; in awake rats, parts of the brain can briefly go offline, resembling the off periods of non-rapid eye movement (non-REM) sleep^56^. Another possibility is that DANs show periods of ‘wake-like’ activity during sleep, because the fly brain can remain active in sleep^15^ as in mammalian REM sleep. Although flies have been reported to have distinct sleep stages associated with altered arousal levels and overall brain activity^30,39^, analysis of individual cell types may help further quantify sleep stages.

We showed that the link between DAN activity and sleep is causal, in line with the previous reports^25,43,45^. Moreover, we observed that DANs also control sensory arousal. Although it may seem intuitive that increased sleep and reduced arousal should always go hand in hand, some circuits and molecules can increase sleep while increasing arousal to stimuli^5,42,57^. We found that silencing the attraction promoting PAM neurons strongly suppresses arousal.

Future studies should elucidate whether DAN-dependent arousal is regulated through downstream MB output neurons, which are also involved in sleep^5,20,22,25,58^. Studies in rodents also report that DAN terminals as well as dopamine D1 receptor-expressing neurons in the reward-processing nucleus accumbens control arousal^59,60^, suggesting that down regulation of attractive dopamine channels is particularly important for animals to stay asleep.

Attractive DANs also played an instrumental role in processing sensory responses during sleep; odor responses were decreased in several attractive PAMs, especially those that we found to encode approach behaviour. In addition, aversive PPL1s increased their responses to further evaluate the odors more aversively. Therefore, during sleep, DANs coordinately tune their responses such that stimuli become less attractive for the animals (Supplementary Fig. 10). Notably, the shift to more negative odor evaluation begins even after shorter periods of rest, possibly as an initial step towards lowering arousal prior to sleep. Sleep is constantly in competition with factors promoting wakefulness such as sexual and hunger instincts^61–63^. Thus, to fall and stay asleep, animals must lower their motivation to pursue attractive stimuli associated with these instincts. Moreover, since sleep is ultimately a vulnerable state, it is likely adaptive to become averse towards stimuli in the surroundings and to take defensive actions upon arousal by dangerous cues. It remains to be determined whether other value assessing systems in the brain exhibit concerted, adaptive changes in valuation of stimuli during sleep.

---

**Supplementary Figure 1.**
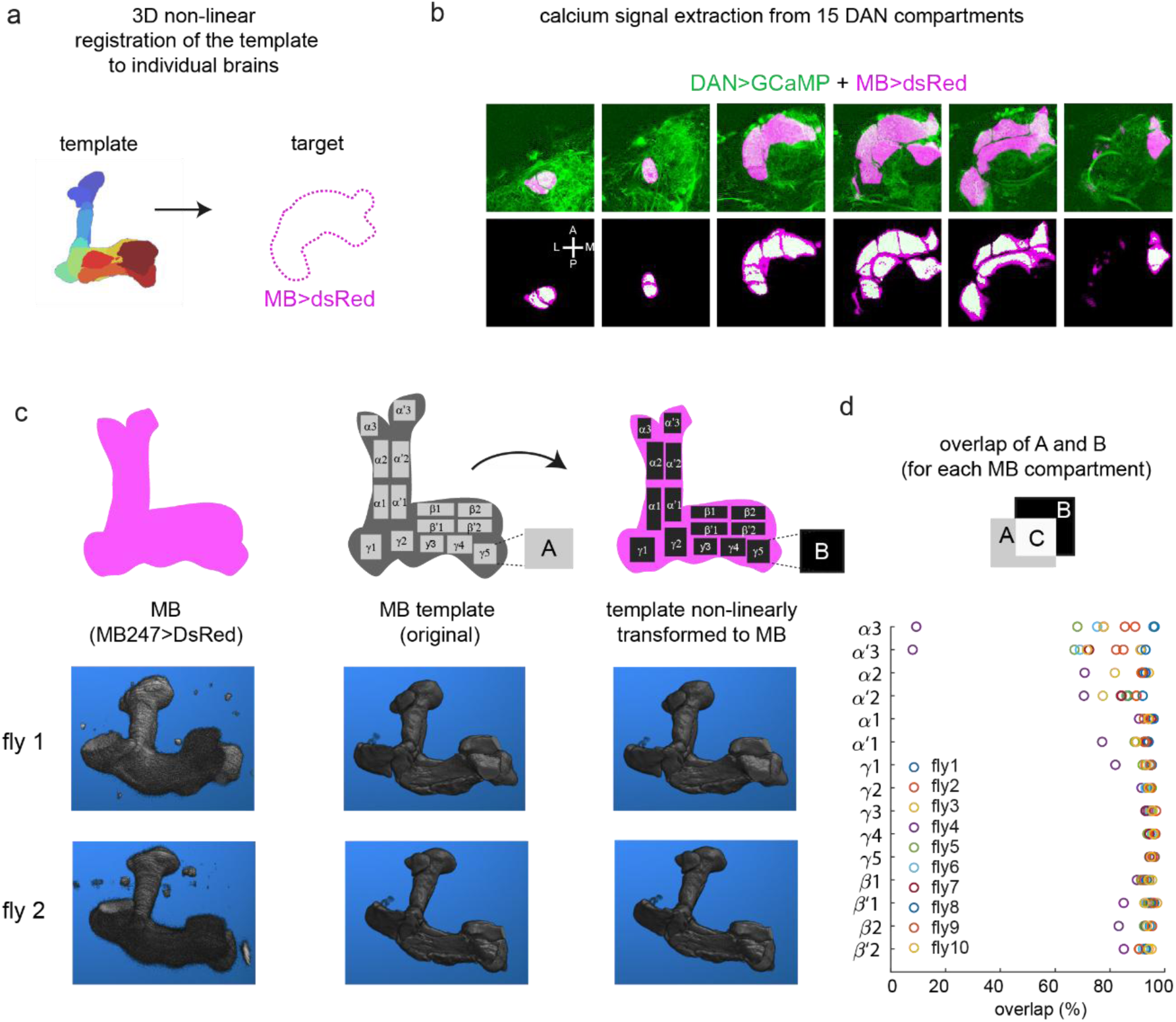
Assessment of quality of registration. (**a**) A template^33^ used to extract fluorescence signals in 15 compartments, which was non-linearly registered to MB247-DsRed signals in individual brains. (**b**) Selected z slice images of MB247-DsRed expression in KCs (magenta) overlayed either with DAN>GCaMP6f expression (green, top) or the template of individual compartments (white, bottom). **(c)** Schema in top panel illustrates how MB compartments were labelled in individual brains, with volumetric images of two example brains underneath. Left panel: volumetric images of MB247-DsRed signals in two different brains. Middle panel: the MB template^33^ was non-linearly registered to the MB247-DsRed signals. Right panel: detection and labelling of compartments in the individual MBs. (**d**) Quality of compartment registration was assessed by calculating the proportion of overlap between each the template compartment “A” and individual labelled compartment “B” (an average was taken from C=A/B and C=B/A, for each compartment). Overlap is shown as a percentage across all brains in which DANs were recorded in Fig.1 and Fig.3.

**Supplementary Figure 2.**
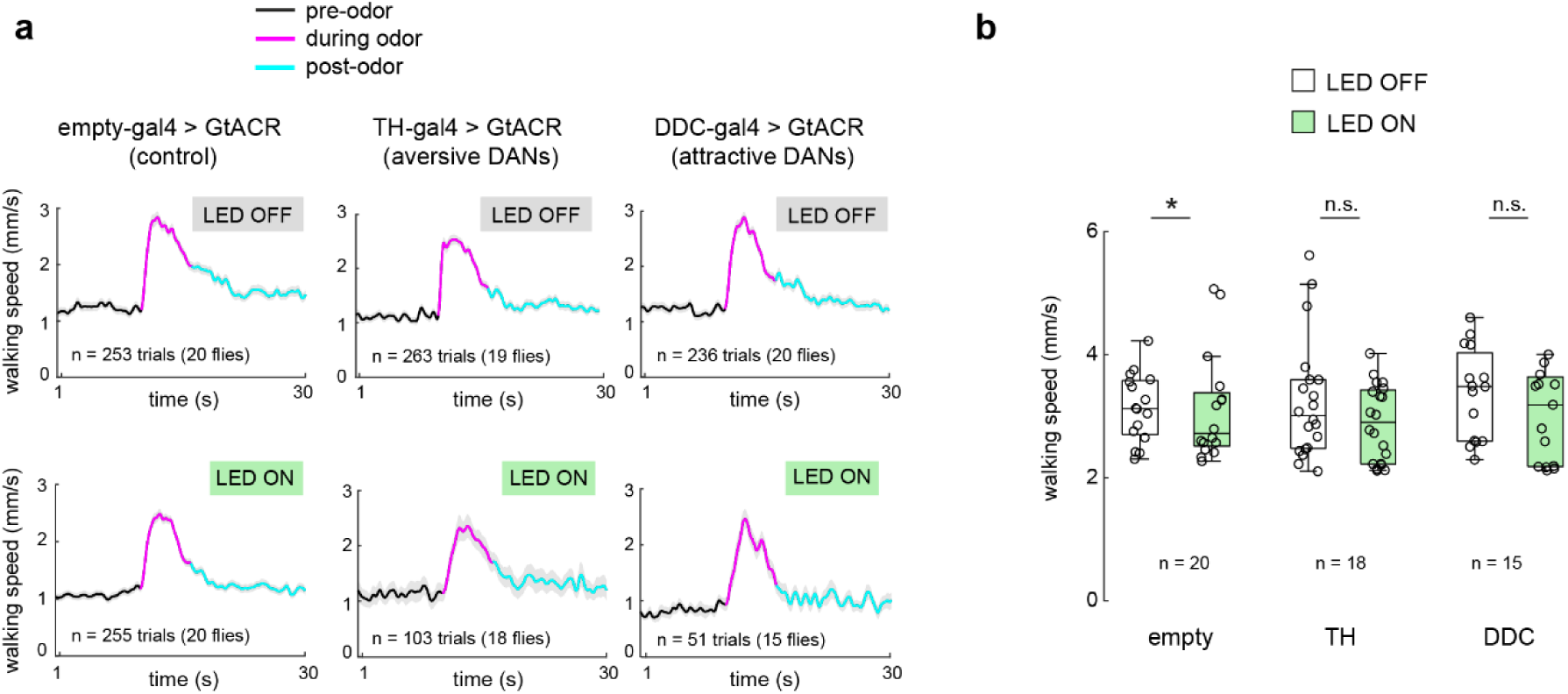
Inhibition of DANs does not impair motor behaviour. (**a**) Average walking speed over time for control and DAN inhibited strains during LED OFF (control) and ON (inhibition) periods, in trials in which flies responded significantly to the odor (with an increased walking speed). Dark line and shaded region represent mean and s.e.m. Number of flies and trials is indicated on graphs. (**b**) Maximum walking speed during odor, with or without LED light (*p = 0.02, Wilcoxon signed-rank test). Boxplots show the median (central line), the interquartile range (box edges), and whiskers (the most extreme datapoints within 1.5 times the interquartile range from the lower and upper quartiles), with each circle showing the average walking speed in a fly. Number of flies is indicated on the graph.

**Supplementary Figure 3.**
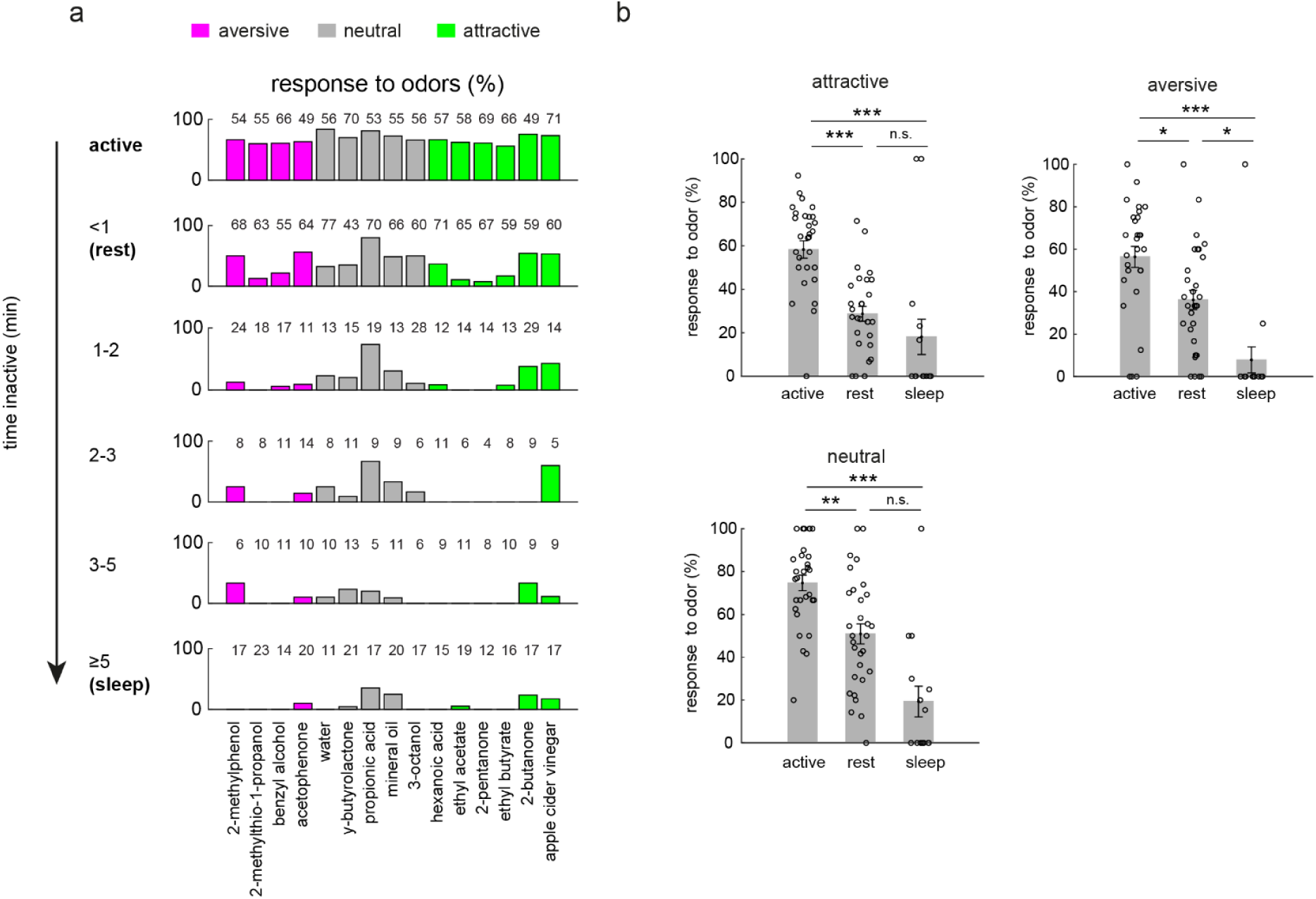
Odor responses to different odors are broadly suppressed during sleep. (**a**) Percentage of trials in which flies responded to each odor in different behavioral states ranging from wake to sleep. Total number of trials pooled across flies for each state is indicated on the graphs. (**b**) The same data as in (**a**) but grouped by aversive, neutral and attractive odors, showing the percentage response of each fly to odors when flies were active, resting or asleep prior to odor onset (mean and s.e.m. across flies are plotted, *p < 0.05, **p < 0.001, ***p < 1e−05, Kruskal-Wallis test with Dunn-Sidak correction for multiple comparisons, n = 30 flies).

**Supplementary Figure 4.**
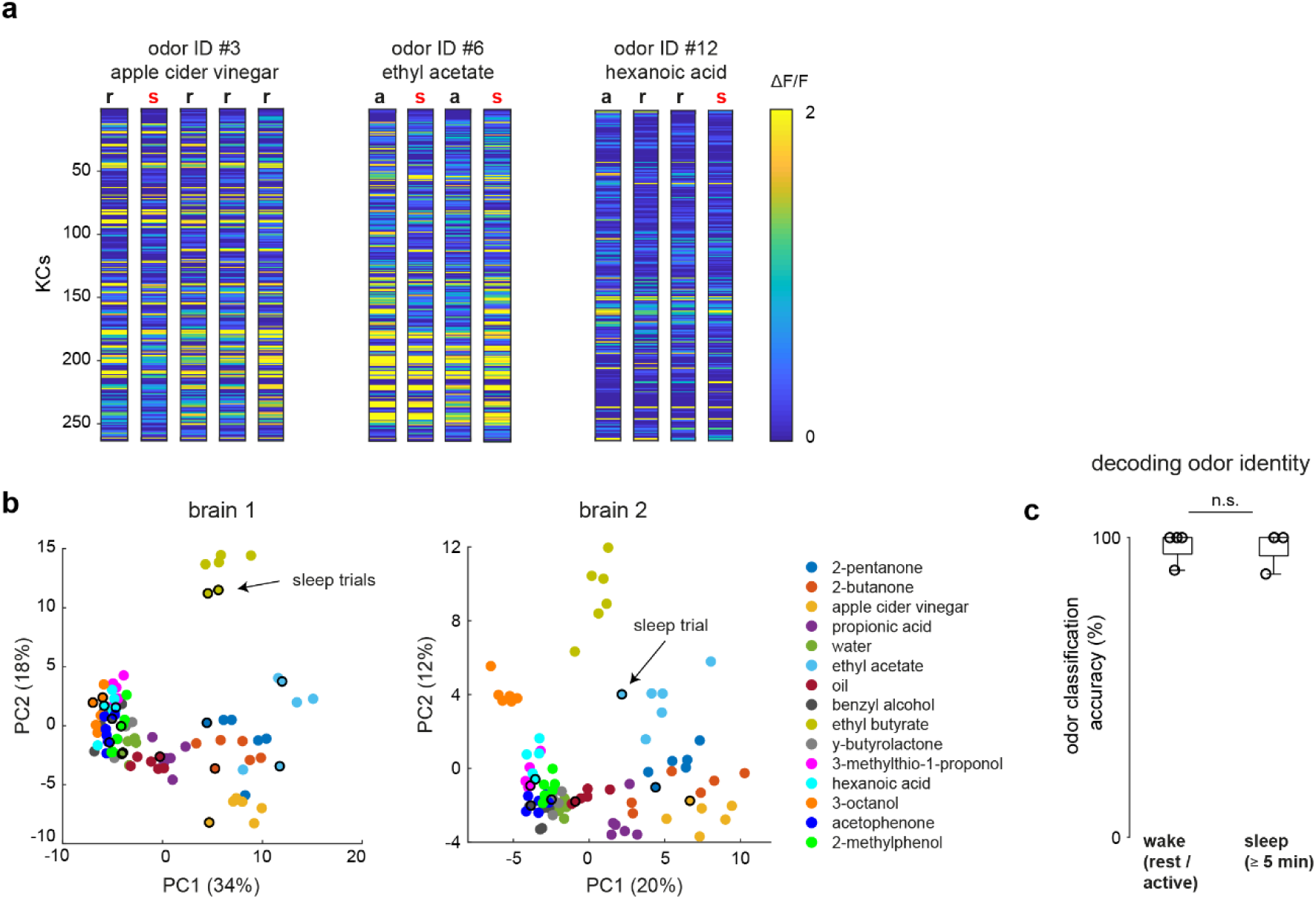
KC odor representations can be accurately decoded during wake and sleep. **(a)** Odor responses (ΔF/F) to 3 different odors when flies were active (‘a’), resting (‘r’) or sleeping (’s’) in an example brain (from Fig. 3d). **(b)** Odor representations in low dimensional space for two example brains (with % variance explained indicated for each PC). Here, each datapoint represents the odor response (ΔF/F) of 263 KCs in brain 1 and 925 KCs in brain 2 (KCs were identified and included in the analysis if they responded significantly to any of the 90 odor trials in the experiment). Odor trials are color coded according to odor identity, from the 15 odors listed in Table S1. Trials in which the fly was asleep are marked with a black outline circle. **(c)** Odor identity decoding using KC responses was accurate during both wake (active or resting) and sleep (n = 17 trials, 4 flies, p = 1.0, Wilcoxon signed-rank test). Boxplot shows the median (central line), the interquartile range (box edges), and whiskers (the most extreme datapoints within 1.5 times the interquartile range from the lower and upper quartiles), with circles showing the accuracy level for each fly.

**Supplementary Figure 5.**
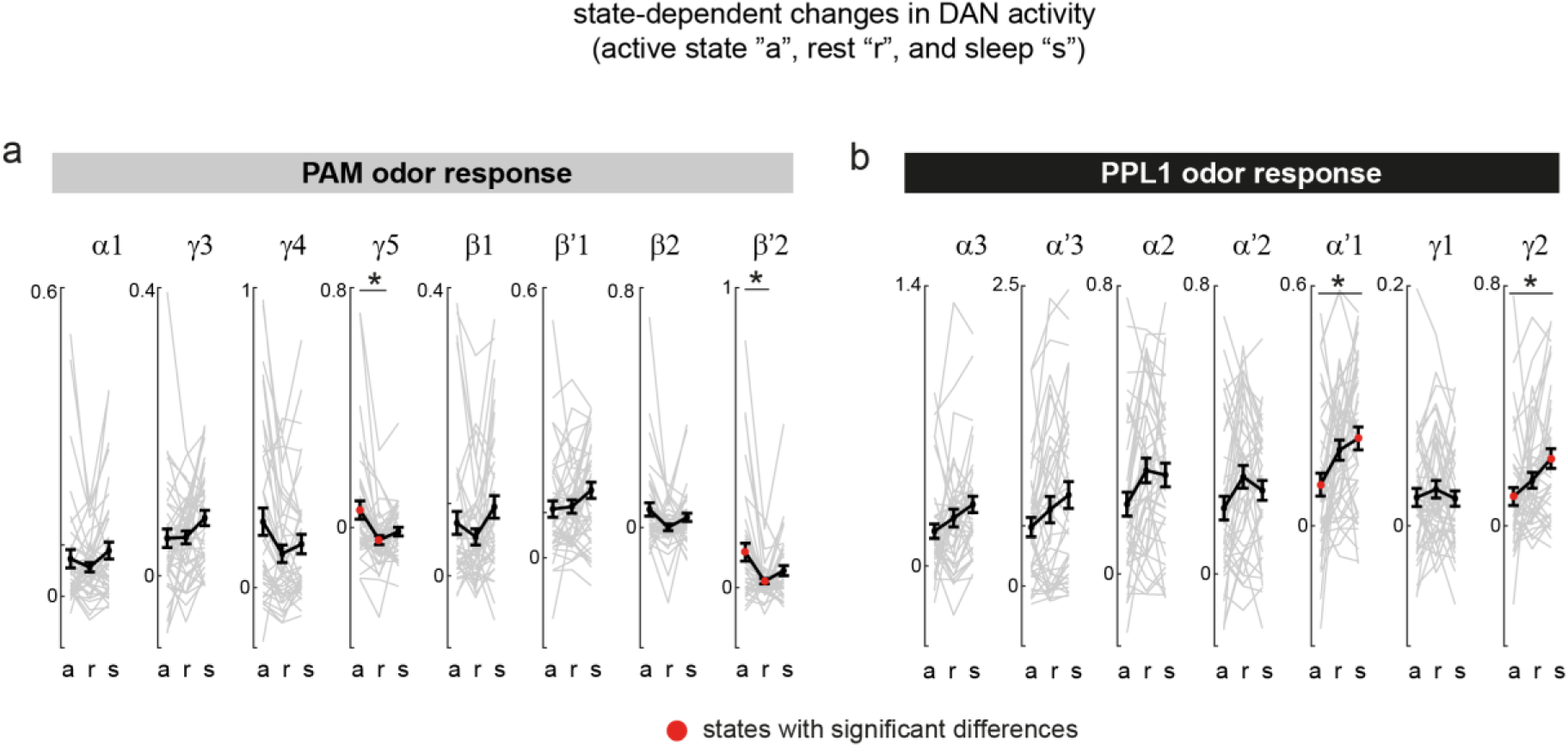
DAN activity during active, rest and sleep states. (**a,b**) Odor responses of DANs in PAM (**a**) and PPL1 (**b**) compartments. Each grey line represents fly-odor matched data across the three behavioural states (a = active, r = rest, s = sleep). Black line and error bar represent mean and s.e.m. States with significant differences are marked in red (n = 41 trials, 10 flies, *p < 0.0011, Wilcoxon signed-rank test, significant after Bonferroni correction for multiple comparisons).

**Supplementary Figure 6.**
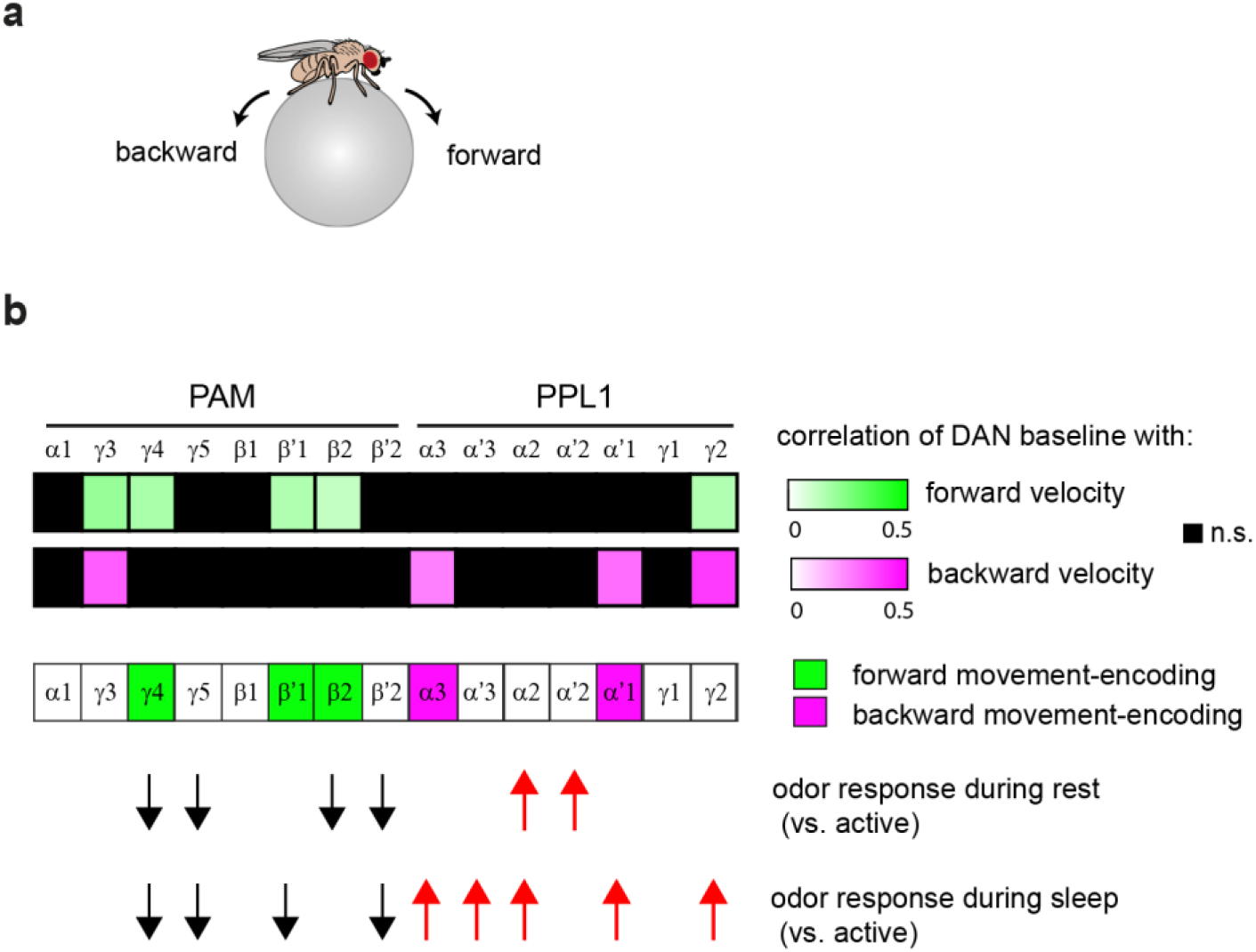
DAN baseline activity is correlated with forward and backward movement during wakefulness. (**a**) Forward and backward walking behaviours. Even when no odor stimulus is present, forward and backward movement may indicate approach-related and avoidance-related behaviours, respectively. (**b**) DAN baseline activity (without odor stimulation) for each compartment was correlated with approach-related behaviours (forward walking) and avoidance-related behaviours (backward walking). Significant correlations are shown as heatmap value of Pearson’s correlation (p < 0.0033, adjusted for multiple comparisons). n = 14 flies (n = 181, 219 and trials for each behaviour type from top to bottom). Below the heatmap, the predominant behaviour encoded by each compartment was colour coded as “approach-encoding” or “avoidance-encoding” based on whether forward or backward walking was correlated selectively and significantly with the DAN baseline activity (where there was no correlation or no selectivity for approach vs avoidance it was left white). Below each compartment, arrows indicate significant changes in odor response that had been observed during rest and sleep (p < 0.0033, Table S2); increased responses occur in avoidance-encoding compartments, whereas reduced responses occur in approach-encoding compartments.

**Supplementary Figure 7.**
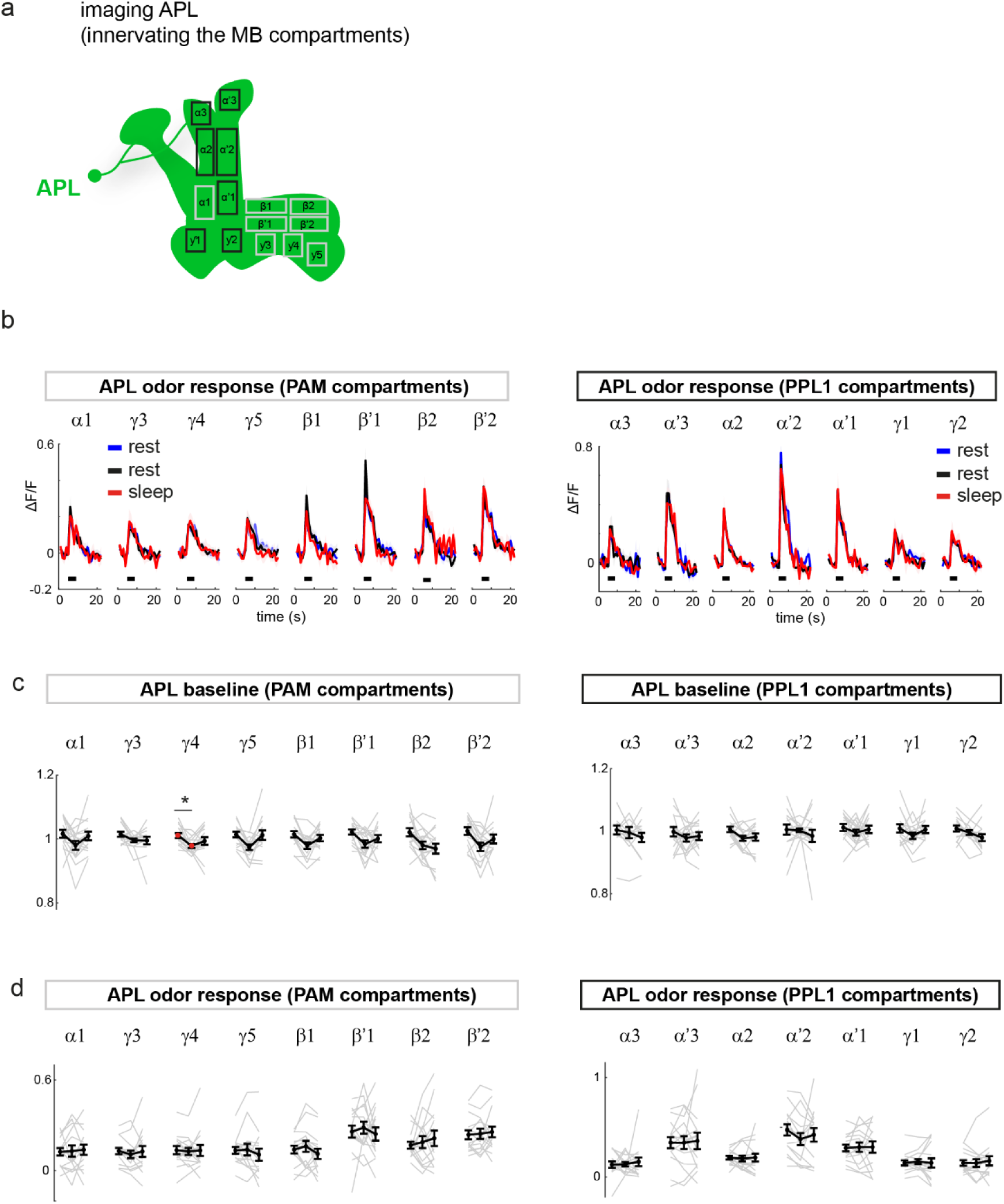
APL activity does not change during rest and sleep. **(a)** Diagram of the APL neuron, which innervates the 15 DAN compartments of the MB. (**b**) Odor responses of APL neurites in each compartment during wakefulness and sleep. Odor application is marked by the grey horizontal bar. (**c,d**) Odor responses (**c**) and baseline activity (**d**) across PAM and PPL1 compartments showed no differences between active (‘a’), rest (‘r’) and sleep (‘s’) states for any compartment, except between active and rest for ƴ4. Each grey line represents fly-odor matched data across the three behavioural states. Black line and error bar represent mean and s.e.m. States with significant differences are marked in red (*p = 0.001, Wilcoxon signed-rank test with Bonferroni correction for multiple comparisons). n = 15 fly-odor matched trials in each state, n = 9 flies.

**Supplementary Figure 8.**
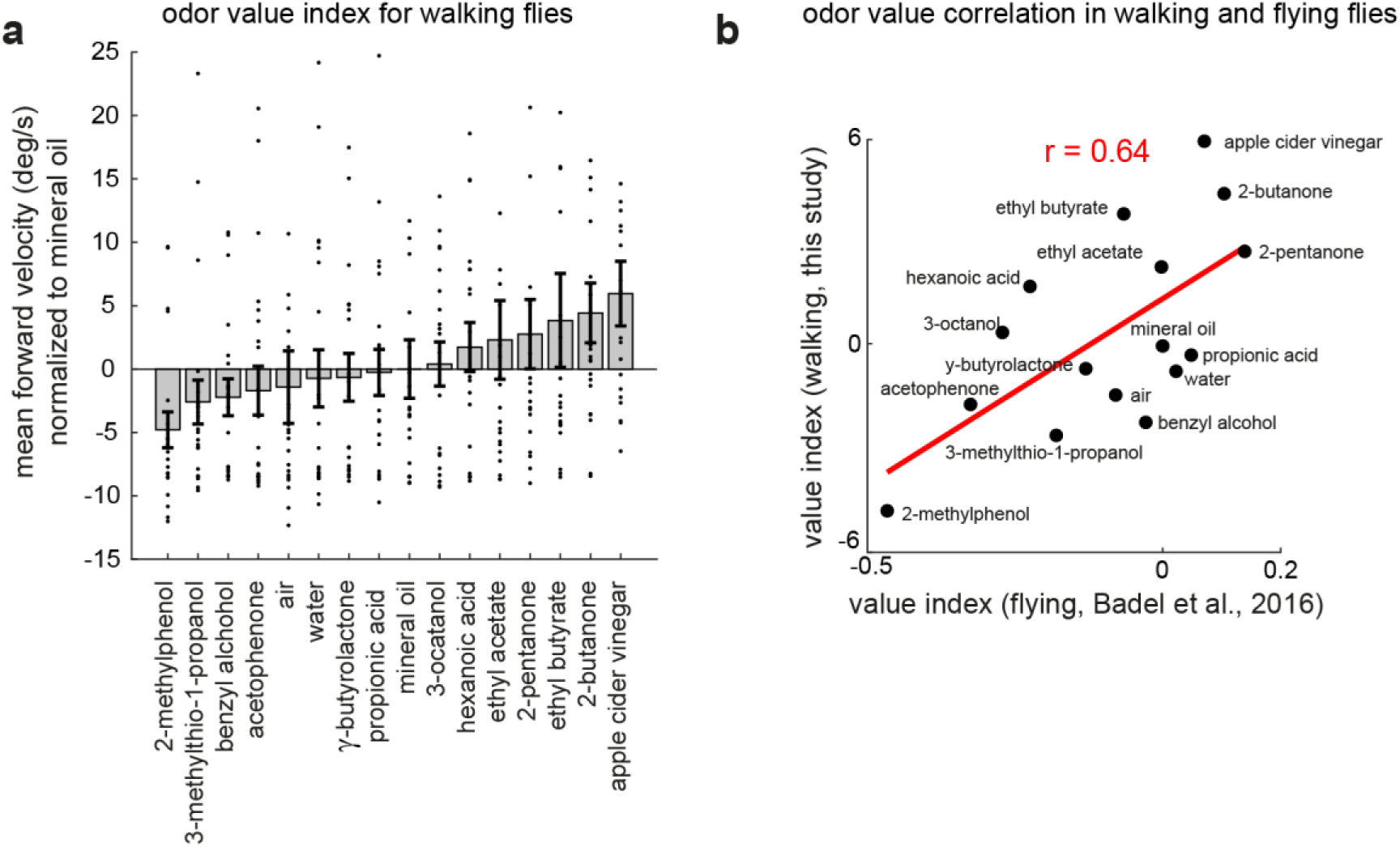
Odor value can be assessed by forward walking behaviour. (**a**) An odor value index for walking flies calculated as the mean forward walking response during the first 3 s of odor application period, subtracted by the value of mineral oil for normalization. Mean and s.e.m across flies are shown, with each dot representing the mean for each fly (n = 6 trials for each odor, n = 22 flies). (**b**) Correlation between the value index in walking flies (**a**) and the value index previously described for flying flies (Badel et al., 2016, Pearson’s correlation = 0.64, p = 0.008).

**Supplementary Figure 9.**
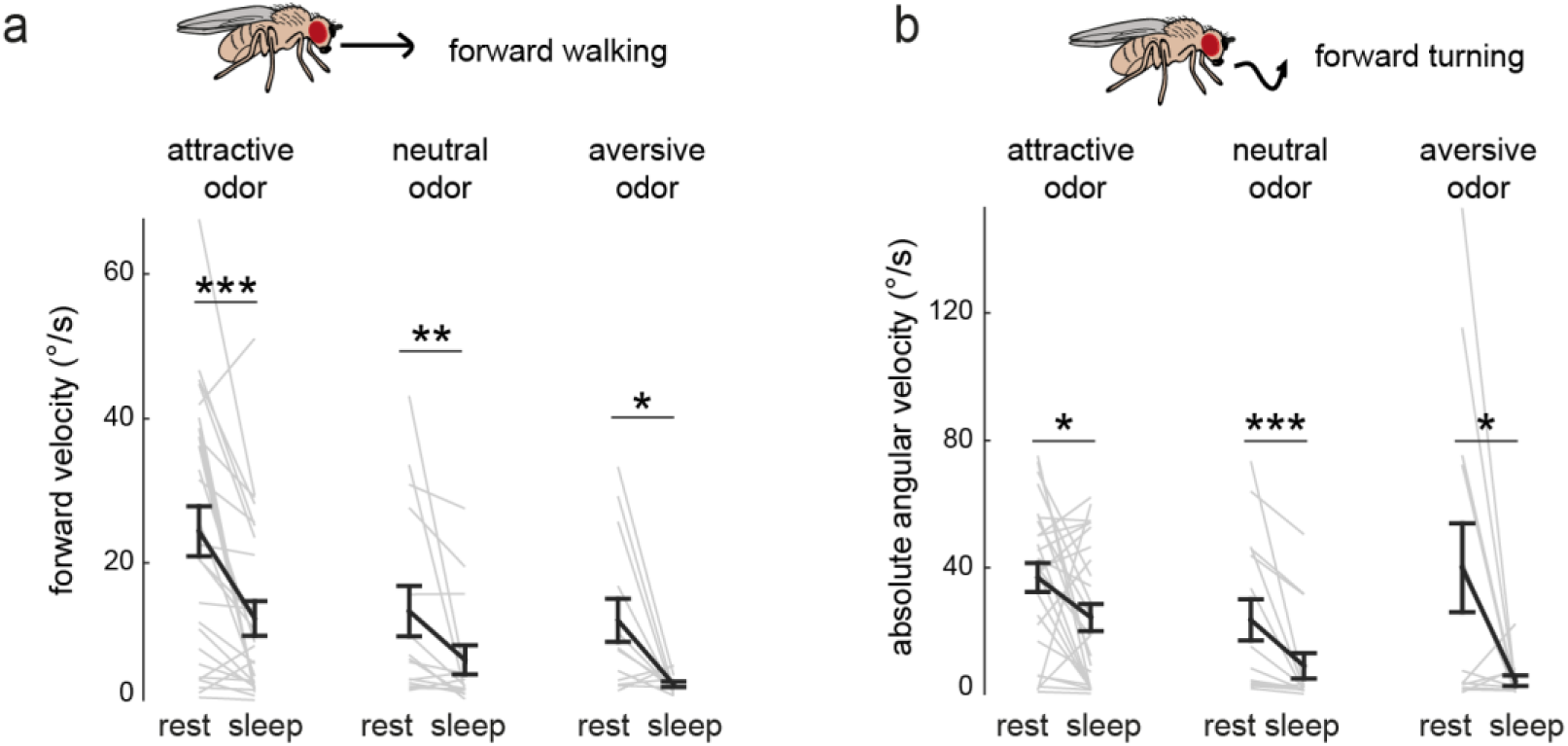
Peak forward and turning velocities in flies awoken from sleep. (**a**,**b**) Maximum values of forward velocity (**a**) and absolute angular velocity when the flies were walking forward during the odor period (**b**) in each trial. Approach behaviours (forward walking and forward turning) were significantly reduced in flies awoken from sleep. Each grey line represents a single trial. n = 49 trials (18 flies) for attractive odors, 36 trials (19 flies) for neutral odors and 32 trials (17 flies) for aversive odors, with all trials matched for fly and odor. Mean and se.m. are indicated by black lines. Wilcoxon signed-rank test was used to compare between rest and sleep, *p <0.05, **p < 0.01, ***p < 0.001.

**Supplementary Fig. 10.**
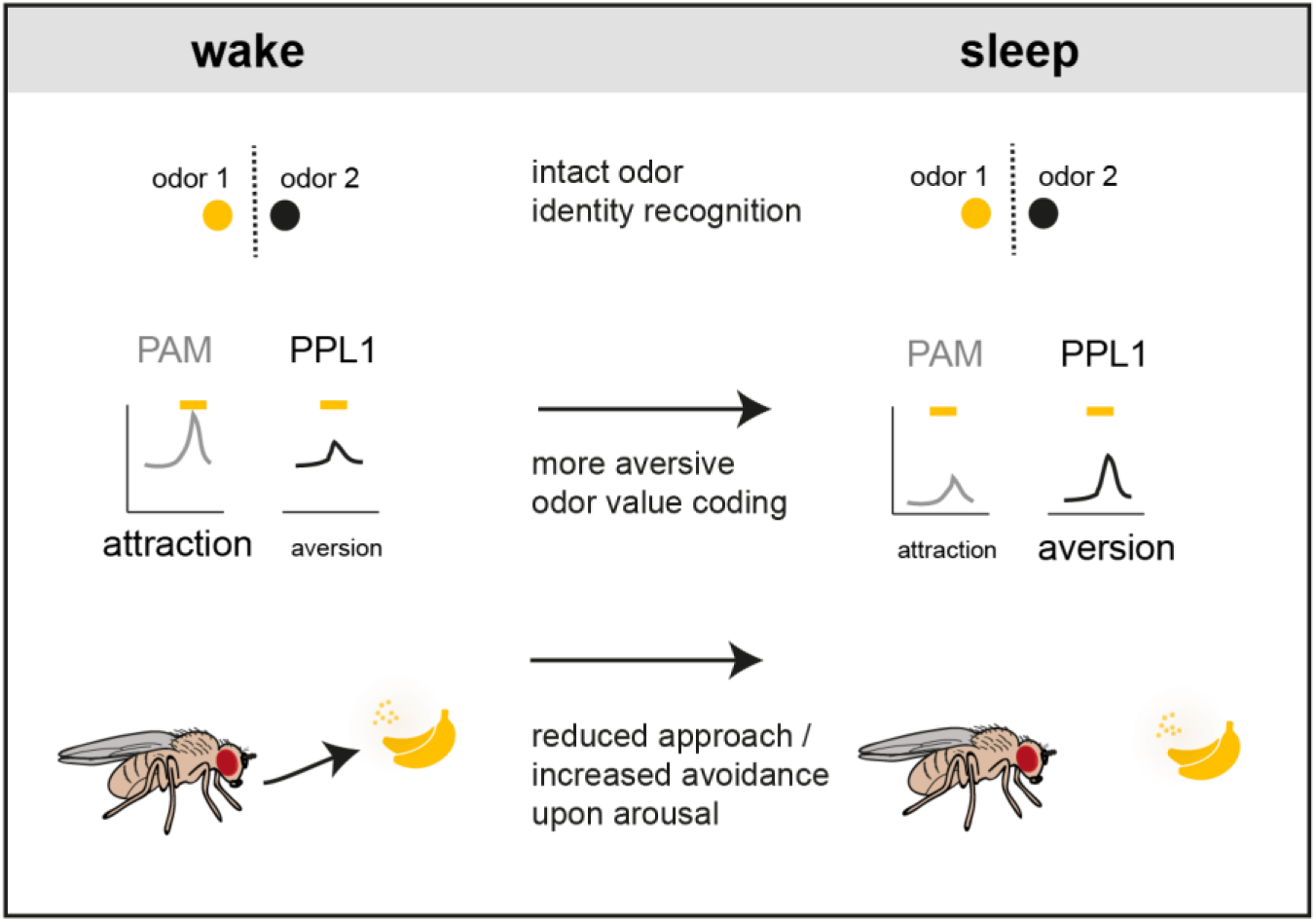
Odor processing and evaluation during wakefulness and sleep. Odor identity recognition remains intact during sleep as KCs can still decode odor identity. However, during sleep, activity of attractive DANs (PAMs) and aversive DANs (PPL1s) changes. Baseline DAN activity is generally lower during sleep, causing lower arousal levels. In contrast, DAN odor responses are stronger in some aversive PPL1s and weaker in some attractive PAMs, together making the odor representation more aversive. Consistently, sleeping flies exhibit reduced attraction and enhanced avoidance behaviour upon arousal to odors. Changes in odor evaluation during sleep may have evolved to prevent arousal and pursuit of attractive odors (such as food, or mates), while maintaining the ability to act defensively when a threatening odor is detected.

**Supplementary Video 1. Simultaneous recording of walking behaviour and KC activity**. Left: A fly walks on an air supported ball, facing a tube delivering the odor (here, apple cider vinegar, an attractive odor). Right: Two-photon Ca^2+^ imaging of all KCs on one side of the brain. A white dot indicates the timing of odor application after which a group of KCs respond with an increase in fluorescence. The video is in real time.

**Supplementary Video 2 (related to Fig. 2). Optogenetic inhibition of dopamine neurons promotes sleep and reduces odor-evoked arousal.** Representative examples of behaviour in flies expressing GtACR in DANs (*TH-Gal4* > *GtACR* and *DDC-Gal4* > *GtACR*) alongside the genetic control (*empty-gal4 > GtACR*), during the LED OFF period and LED ON (inhibition) period. During the “LED ON” period, trials in which *TH-Gal4* > *GtACR* and *DDC-Gal4* > *GtACR* flies had been inactive for at least 5 min are displayed. These flies remained inactive and unresponsive to odors. In contrast, *empty-gal4 > GtACR* control flies remained awake with frequent bouts of walking and responses to odors. The odor delivered was apple cider vinegar for all flies and trials in this video.

**Table S1.**
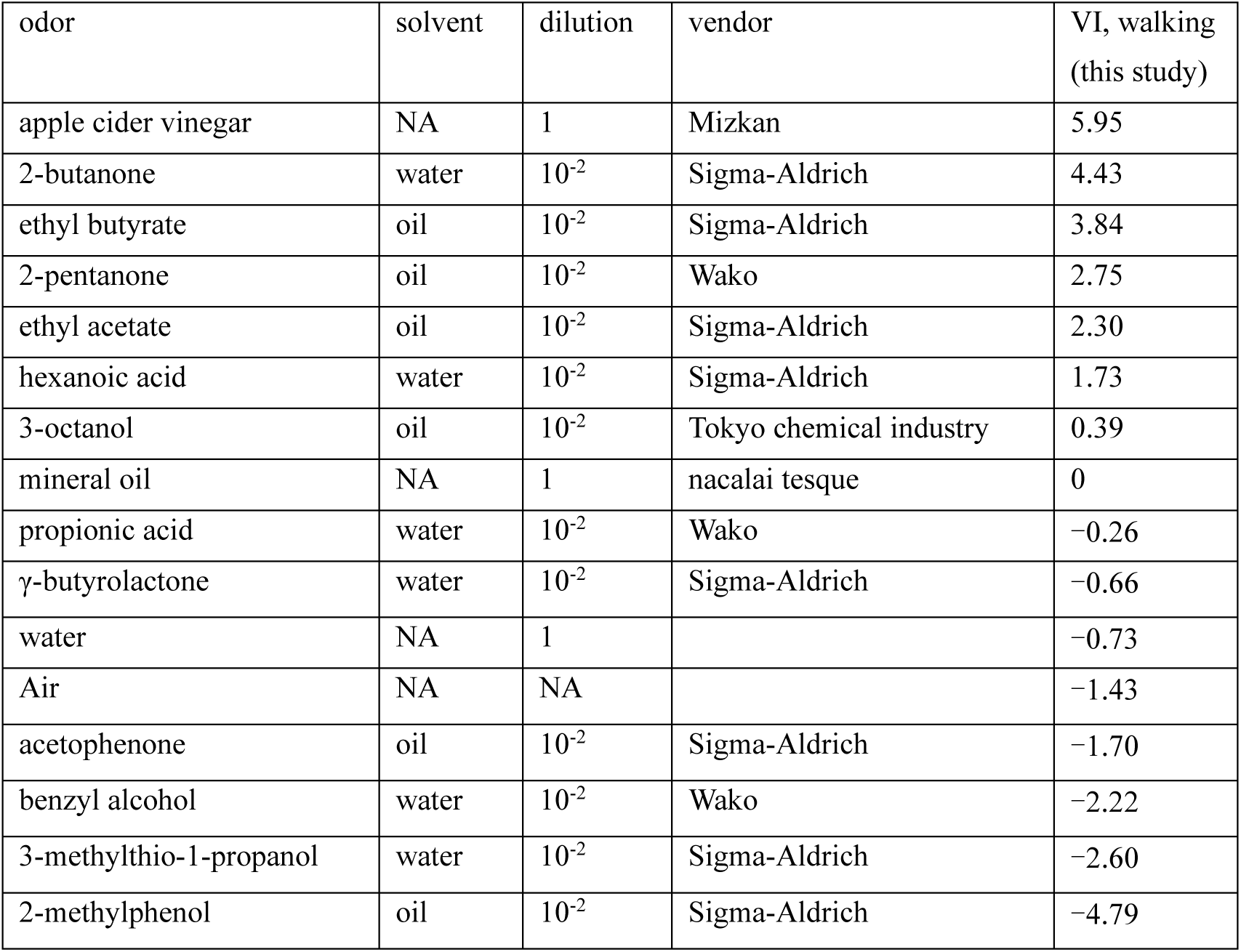
List of odors used in this study.

| odor | solvent | dilution | vendor | VI, walking<br>(this study) |
| --- | --- | --- | --- | --- |
| apple cider vinegar | NA | 1 | Mizkan | 5.95 |
| 2-butanone | water | $10^{-2}$ | Sigma-Aldrich | 4.43 |
| ethyl butyrate | oil | $10^{-2}$ | Sigma-Aldrich | 3.84 |
| 2-pentanone | oil | $10^{-2}$ | Wako | 2.75 |
| ethyl acetate | oil | $10^{-2}$ | Sigma-Aldrich | 2.30 |
| hexanoic acid | water | $10^{-2}$ | Sigma-Aldrich | 1.73 |
| 3-octanol | oil | $10^{-2}$ | Tokyo chemical industry | 0.39 |
| mineral oil | NA | 1 | nacalai tesque | 0 |
| propionic acid | water | $10^{-2}$ | Wako | -0.26 |
| γ-butyrolactone | water | $10^{-2}$ | Sigma-Aldrich | -0.66 |
| water | NA | 1 |  | -0.73 |
| Air | NA | NA |  | -1.43 |
| acetophenone | oil | $10^{-2}$ | Sigma-Aldrich | -1.70 |
| benzyl alcohol | water | $10^{-2}$ | Wako | -2.22 |
| 3-methylthio-1-propanol | water | $10^{-2}$ | Sigma-Aldrich | -2.60 |
| 2-methylphenol | oil | $10^{-2}$ | Sigma-Aldrich | -4.79 |

**Table S2.** Statistical comparison of DAN odor responses (related to Supplementary Fig. 6). The results of statistical tests comparing the DAN odor responses during different behavioural states. Numbers indicated p values and arrows indicate direction of change compared to the active state. n = 76 fly-odor matched trials for each state. *p < 0.0033 with Wilcoxin sign-rank test (Bonferroni correction for multiple comparisons).

|  | Rest (vs active) | Sleep (vs active) |
| --- | --- | --- |
| $\alpha 1$ (PAM) | 0.15 | 0.63 |
| $\gamma 3$ (PAM) | 0.19 | 0.15 |
| $\gamma 4$ (PAM) | 7.1e-07 * ↓ | 3.9e-05 * ↓ |
| $\gamma 5$ (PAM) | 2.6e-08 * ↓ | 9.8e-06 * ↓ |
| $\beta 1$ (PAM) | 0.06 | 0.14 |
| $\beta'1$ (PAM) | 0.52 | 0.002 * ↑ |
| $\beta 2$ (PAM) | 3.0e-05 * ↓ | 0.007 |
| $\beta'2$ (PAM) | 6.9e-08 * ↓ | 2.86e-06 * ↓ |
| $\alpha 3$ (PPL1) | 0.06 | 3.8e-06 * ↑ |
| $\alpha'3$ (PPL1) | 0.11 | 0.001 * ↑ |
| $\alpha 2$ (PPL1) | 1.2e-05 * ↑ | 0.001 * ↑ |
| $\alpha'2$ (PPL1) | 7.8e-05 * ↑ | 0.06 |
| $\alpha'1$ (PPL1) | 0.004 | 9.3e-05 * ↑ |
| $\gamma 1$ (PPL1) | 0.80 | 0.99 |
| $\gamma 2$ (PPL1) | 0.14 | 4.6e-05 * ↑ |

## Methods

### Fly Stocks and husbandry

Flies were raised on conventional cornmeal agar medium at 25 °C in a 12 h light, 12 h dark cycle. Post-eclosion, adult virgin females were collected and maintained at a density of ∼10 flies per vial, with the day night cycle reversed (9 am - 9 pm lights off), at 23 °C to acclimatize to the temperature of the experimental room. All experiments were performed on virgin adult females at 3 - 7 days post-eclosion. For optogenetic experiments, flies were fed food containing 0.2 mM all-trans-retinal (Sigma Aldrich, R2500) for at least 2 days prior to the experiment.

Fly stocks used in this study were: *R13F02-Gal4(attP2)*^1^ (Bloomington # 48571), *MB247-DsRed*^2^ (gift from André Fiala), *R58E02-LexA(attP40)* (Bloomington # 52740), *LexAop-GCaMP6s(attP1)* (Bloomington # 44274), *UAS-opGCaMP6s(attP5)*^3^ (gift from G. Rubin), *20XUAS-IVS-Syn21-opGCaMP6f-p10(su(Hw)attP8)*^3^ (Janelia Research Campus), *UAS-mCherry.NLS* (Bloomington # 38425), *UAS-GtACR1-EYFP*(attP2)^4^ (gift from Adam Claridge-Chang), *TH-LexA*^5^ (gift from Hiromu Tanimoto), *tubPGal80ts* (Bloomington #7019), *empty-Gal4* (Bloomington #68384), *DDC-Gal4*^6^ (Bloomington #7009), *TH-Gal4*^7^ (gift from Hiromu Tanimoto), *VT43924-Gal4.2*^8^ (gift from Andrew Lin).

Fly genotypes used in this study were as follows:

Figure 1, Figure 3, Supplementary Figures1, 3, 4, 5, 6, 8
*20XUAS-IVS-Syn21-opGCaMP6f-p10(su(Hw)attP8);;R13F02-Gal4*
*R58E02-LexA, MB247-DsRed; TH-LexA, LexAop-GCaMP6s / LexAop-GCaMP6s*
*R58E02-LexA, MB247-DsRed; TH-LexA, LexAop-GCaMP6s / TM2*

Figure 2, Supplementary Figure 2
*UAS-GtACR1-EYFP(attp2)/+; tubP-Gal80ts/+; TH-Gal4/+*
*UAS-GtACR1-EYFP(attp2)/+; tubP-Gal80ts/+; DDC-Gal4 /+*
*UAS-GtACR1-EYFP(attp2)/+; tubP-Gal80ts/+; empty-Gal4/+*
*TH-Gal4* and *DDC-Gal4* had been backcrossed for six generations to wild type established in the Michael Dickinson’s laboratory^9^.

Figure 4, Supplementary Figure 9
*20XUAS-IVS-Syn21-opGCaMP6f-p10(su(Hw)attP8);;R13F02-Gal4*
*R58E02-LexA, MB247-DsRed; TH-LexA, LexAop-GCaMP6s / LexAop -GCaMP6s*
*UAS-opGCaMP6s(attP5), UAS-mCherry.NLS; R13F02-Gal4(attP2)*
*20XUAS-IVS-Syn21-opGCaMP6f-p10(su(Hw)attP8);;R13F02-Gal4/R23E10-Gal4*

Supplementary Figure 7
*20XUAS-IVS-Syn21-opGCaMP6f-p10(su(Hw)attP8);* MB247-DsRed/+*;VT43024-gal4.2/+*

### Odorants

2-pentanone (Cat#: 133–03743; CAS: 107-87-9, Wako), 2-butanone (Cat#: 04380; CAS: 78-93-3, Sigma-Aldrich), Apple cider vinegar (Mizkan), Propionic acid (Cat#: 163–04726 CAS: 79-09-4, Wako), mineral oil (nacalai tesque, 2334-85), Ethyl acetate (Cat#: 319902; CAS: 141-78-6, nacalai tesque), Benzyl alcohol (Cat#: 027–01276; CAS: 100-51-6, Wako), Hexanoic acid (Cat#: 153745; CAS: 142-62-1, Sigma-Aldrich), Ethyl butyrate (Cat#: E15701; CAS: 105-54-4, Sigma-Aldrich), γ-butyrolactone (Cat#: B103608; CAS: 96-48-0, Sigma-Aldrich), 3-methylthio-1-propanol (Cat#: 318396; CAS: 505-10-2, Sigma-Aldrich), 3-octanol (Cat#: O0121; CAS: 589-98-0, Tokyo Chemical Industry), Acetophenone (Cat#: A10701; CAS: 98-86-2, Sigma-Aldrich), 2-methylphenol (o-cresol) (Cat#: C85700; CAS: 95-48-7, Sigma-Aldrich).

### Olfactory Stimulation

Odors were diluted with mineral oil (nacalai tesque, 2334-85) or water except for apple cider vinegar, which was not diluted. Odors were delivered as described previously^10^, using a multi-channel olfactometer controlled by custom MATLAB (MathWorks) code via a data acquisition interface (NI cDAQ-9178, NI9264 National Instruments). Briefly, an air flow (250 mL/min) passed through 4 mL of odor solution in a vial converged with a main air steam (1550 mL/min), after which a small portion of it was directed frontally to the fly through a ø2 mm outlet tube placed approximately 10 mm away from the fly’s antennae. The speed of the odorized air remained constant throughout the experiment and during odor delivery at 0.3 m/s. Odors used in this study are listed in Table S1. All odors were applied for 4 s (or 6 s, in DAN experiments) with an inter-trial interval of 60 s. Experiments consisted of 6 blocks of 15 odors (plus an air control) for the KC and DAN recordings with the order of odor application randomized for each block. For experiments that did not require decoding, fewer odors were used to obtain more trials of matched odors during sleep and wake: 20 blocks of 3 odors for APL imaging, and 10 blocks of 6 odors (2 attractive, 2 neutral, and 2 aversive) for optogenetic inhibition experiments. Stimulus timing onset was defined as the time the odor reached the fly, which was measured previously^10^.

### Fly preparation

To increase the probability of flies falling asleep during experiments, flies were socialized (housed at a density of ∼ 10 flies per vial), as socialization increases sleep in *Drosophila* ^11^. Some flies were sleep deprived for 24 h prior to the experiment using a gentle 1 s vibration delivered at random intervals centered around 15 s. The vibration was provided with an LC4 controller (Trikinetics) using a shaker (SIBATA, 050630-37) set at a level just strong enough to dislodge flies at the top of the vial. Food was placed at the bottom of the vial so that flies had continuous access to food during sleep deprivation.

To tether the fly to a custom recording plate, individual animals were briefly cold-anesthetized on ice (∼30 s) and placed in a holder on a Peltier plate kept at 4 °C. The fly head and the upper edge of the thorax were fixed to the recording plate using ultraviolet-curing adhesive (NOA63, Norland). For KC experiments, the head was angled down in this step to view more posterior side of the head as described previously ^12^. The proboscis was glued in a retracted position to prevent brain movement during the recordings, and the body position was adjusted to let the fly walk smoothly when placed on the air-supported ball. After immersing the head in saline solution (103 mM NaCl, 3 mM KCl, 5 mM N-tris (hydroxymethyl) methyl-2-aminoethane-sulfonic acid, 8 mM trehalose, 10 mM glucose, 26 mM NaHCO_3_, 1 mM NaH_2_PO_4_, 1.5 mM CaCl_2_, and 4 mM MgCl_2_, osmolarity adjusted to 270-275 mOsm), a portion of the head cuticle was removed, along with trachea and fat bodies that obscured the imaging region. The muscle of the frontal pulsatile organ (muscle 16) was removed to prevent brain movement. During the experiment, saline was bubbled with 95% O_2_/5% CO_2_ at a rate of 2 mL/min.

### *In vivo* two-photon Ca^2+^ imaging and behavioural recordings

Flies were placed on an air-supported ball (PP ball, 6 mm in diameter, Spherotech GmbH, Germany), floating in a custom-built ball holder (Electronics workshop, Zoological Institute, University of Cologne). The airflow supporting the ball was adjusted to the minimal level that would spin the ball when the fly stepped on it. The ball and the fly were backlit by an infrared light (730 nm, M730L4, Thorlabs) passing through a piece of white polyacetal diffusion material (Kureha Extron). The ball was painted with a black, oil-based pen (Name Pen, Sakura Color Products Corporation, Japan). A camera (Firefly, FMVU-03MTC-CS, FLIR) with a 730 nm bandpass filter covering the lens (FL730-10, Thorlabs) provided a sideview of the fly on the ball, and was used to collect videos at 30 fps with Fictrac software ^13^. Prior to each experiment, the fly’s position was adjusted until it could smoothly walk on the ball.

Fluorescence signals were acquired using a two-photon microscope (LSM 880 with Airyscan Fast, Zeiss) equipped with a piezo motor (P-725.2CD PIFOC, PI) that drives a water immersion objective lens (W Plan-Apochromat, 20x, numerical aperture 1.0) along the z axis. GCaMP was excited at 930 nm with a Ti:Sapphire laser (MaiTai eHP DeepSee, Spectra-Physics). GCaMP signal was collected with an internal 32-channel GaAsP detector through a 420-480 + 495-550 bandpass filter.

The KC population was imaged in FAST mode by scanning a 133.12 x 133.12 x 67.2 µm volume with a pixel size of 0.32 x 0.32 µm and a z slice interval of 1.92 µm, resulting in a scanning rate of 1.8 s/volume.

The DAN population was imaged in LSM mode by scanning a 103.68 × 103.68 x 170 µm volume with a pixel size of 1.62 x 1.62 µm and a z slice interval of 5 µm, resulting in a scanning rate of 1.228 s/volume. After Ca^2+^imaging, high-resolution images of DsRed and GCaMP were taken at 0.405 x 0.405 µm resolution, with a z slice interval of 1 µm. DsRed was excited with a 543 nm CW laser and the fluorescence was detected through a 495-535+LP555 filter emission filter. In cases where brain movement was observed, 1 µM tetrodotoxin (TTX, Cat# 32775-51, nacalai tesque) was bath-applied for 10 min prior to taking the high-resolution images.

### Optogenetic Manipulation

For behavioural experiments involving DAN silencing through the activation of GtACR, flies were prepared in the same way as for two-photon imaging (with cuticle dissected, brain exposed and fresh saline applied), except that the proboscis was not glued. Removing the cuticle allowed us to use a lower level of green light to optically inhibit DANs. A 530 nm LED (M530F2, Thorlabs) was directed to the brain from above, via an optic fiber (M118L03, Thorlabs) placed at an angle of 30°, ∼0.5 cm from the brain. LED pulses were delivered at two light levels, 3.8 and 8.5 μW/mm^2^, at a frequency of 50 Hz (repetition of a square wave with 10 ms duration). Arousal to odors was monitored by exposing flies to 2 aversive odors (3-octanol and 2-methylphenol), 2 neutral odors (propionic acid and y-butyrolactone), and 2 attractive odors (2-pentanone and apple cider vinegar), with 60 s inter-stimulus interval. Each odor was repeated across two sets of two contiguous blocks in a different order for each block, with block sets alternating between green light ON and OFF. The order of light ON and OFF was reversed in half of the experiments to balance out block order effects. Sleep was quantified by calculating the percentage of time that each fly spent asleep. Arousal was quantified by calculating the percentage of trials in which each fly responded significantly (above the walking speed threshold) during the odor application period.

### Data analysis

Image processing and analyses were performed using Fiji ^14^ (version 1.53t), ANTs (Advanced Normalization Tools) version 2.4.2.post31-g74afcc5 ^15,16^, and custom code written in MATLAB (MathWorks, versions 2013a, 2019a and 2024a).

#### Image registration and signal extraction

Brain motion during Ca^2+^ imaging was corrected by applying 3D rigid registration (to remove the effects of slow drift of the brain through the experiment) and 2D, XY plane-by-plane rigid registration (to remove the effects of high-frequency brain movements). For 2D registration, each XY planar image at each time frame was registered at sub-pixel resolution to a template, which is the mean baseline image of a trial that least deviated from all trials in the experiment. This was done by shifting the planar image along X, Y, and Z axes to maximize the correlation between each image and the template.

Analysis of KC images was in principle conducted as described previously ^12^. Volumes of Interest (VOIs) corresponding to KCs were extracted and included in the analyses when they showed a significant response to any odor in any trial during the experiment, using a 2-sample t-test to compare fluorescence during baseline and odor periods (4 time frames each). This resulted in analysis of 1652 responding KCs in total across all trials in Fig. 3h rest vs sleep (263, 944, 248, 197 cells in 4 flies), 1718 responding KCs in total across all trials in Fig. 3h rest vs active (263, 247, 338, 248, 197, 425 cells in 6 flies), and 1829 responding KCs in total across all trials in Supplementary Fig. 4c (263, 944, 197, 425 cells in 4 flies).

Analysis of DAN and APL images was conducted as described previously ^17^. This involved 3 steps: 1) The 3D template volume of the 15 MB compartments (JFRC2013 standard brain ^18^) was registered non-linearly to a high resolution MB247-DsRed signal within the MB of each fly brain, creating a high resolution master image for each brain, 2) functional GCaMP images were registered to the master image and 3) GCaMP signal was extracted for each compartment using the registered template. The percent overlap of the MB image to the template was calculated following registration, and brains were discarded if the overlap was below 78% (most brains achieved above 90% overlap) or if fluorescence signals were not detected in each of the 15 compartments.

#### Analysis of Calcium imaging data

Odor responses were quantified as ΔF/F ((F-F_baseline_)/F_baseline_) where F_baseline_ is an average fluorescence across 5 s preceding the stimulus onset. To compare odor responses between behavioural states while controlling for variation between different odors and different flies, responses to a particular odor were included in the analysis only when they were collected in each of the behavioural states being compared in the same fly. In other words, an odor was not included in the analysis if all the responses happened to be collected exclusively in active, rest, or sleep states in a fly (no comparison can be made between states for this odor). When a fly had multiple trials for the same odor in the same behavioural state, the average response was taken (an average was not taken for decoding analysis. See below). Baseline activity was compared between states using the same datasets. Baseline activity for each trial was quantified as an average activity over 4 time frames prior to odor application (F_baseline_), normalized to the overall average baseline across all trials within a block for each fly (F_baselineOverall_). This procedure was introduced to correct for the long-term drift in fluorescence across multiple blocks, which may interfere with the detection of differences in F_baseline_ between states, using the following equation:

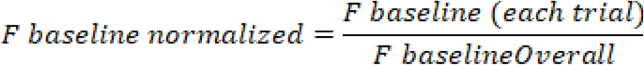

For APL experiments, which had shorter blocks consisting of only 3 odor trials, and for which the baseline remained stable throughout the experiment, F_baselineOverall_ was calculated as the average F_baseline_ across all the trials throughout the experiment for each fly.

#### Decoding of odor identity, odor value and behavioural state

For decoding of odor identity from KC responses, we built a model for each fly, as the identity of KCs is unique for each fly (flies were excluded from analysis when they did not show sufficient sleep trials to make comparisons between behavioural states). To reduce noise in the data, principal component analysis was applied to the population KC responses. In all three analyses, we used the first eight principal components, accounting for, 87.57 ± 2.96% (Fig. 3h, rest vs 3-min sleep), 89.32 ± 4.19% (Fig. 3h, rest vs active), and 79.29 ± 7.49% (Supplementary Fig. 4c) of the total variance, respectively. The decoding models for each fly were built using linear discriminant analysis, which linearly separates different odor identities. At least 2 trials from 3-15 odors were used for training. The same number of trials from the same odors for each behavioural state were withheld for testing. Decoding for each fly was repeated 100 times with different selections of training and test trials. Accuracy for each fly was calculated as the proportion of trials in which the odor identity was correctly predicted.

Decoding of odor value from DAN responses was conducted in three steps: quantifying the odor value, preprocessing and classifying the DAN responses to training and test data, and building a decoding model.

To quantify the odor value for walking flies, a value index that ranks odors from attractive to aversive was created based on odor approach behaviour (forward walking) (Supplementary Fig. 8a, Table S1). The value of each odor was calculated as the mean forward velocity during the first 3 s of odor application period, normalized to mineral oil (by subtracting the mean value of mineral oil) (Supplementary Fig. 8a). Attractive odors were defined as those with an odor value above 1.5, and aversive odors were defined as those with a value below − 1.5, with neutral odors in between. To calculate odor values, we restricted our analysis to trials in which flies were awake at the time of odor delivery (active or rest) and where they responded significantly to the odor (with a significant increase in walking speed, comparing frames in the 6 s baseline period before the odor to 6 s period during the odor application).

DAN odor responses (ΔF/F) in each MB compartment were used for predicting odor value. Each trial was indexed as active, rest or sleep based on the average walking speed prior to and during the odor delivery. The same number (93) of odor-fly matched active, sleep and rest trials were withheld as test data. The remaining data (217 trials in the rest state) was used for training the model.

The decoding model was built using partial least squares (PLS) regression, a method known to be efficient in cases with many predictors and robust to collinearity in the predictors ^19^. First, the number of PLS components was optimized by running PLS regression with 10-fold cross validation on the training data set, resulting in the identification of 5 PLS components as giving the highest accuracy for predicting odor value. Next, the actual model was built using the entire training data set, following which odor value was predicted for withheld active, rest and sleep test trials. This was repeated 100 times with random sampling of training and test data, and the mean predicted values were calculated as an average across the 100 models. The accuracy of decoding was quantified with the prediction R^2^.

For decoding of behavioural states from DAN baseline activity, a quadratic discriminant decoder was used to discriminate the 3 states, using either baseline from the 8 PAM or 7 PPL1 compartments. Data from 11 flies that slept at least once during the experiment were used for the analysis (non-sleeping flies were excluded). Baseline activity was processed as described earlier for each fly after which the data were pooled across flies for each compartment. To reduce noise in the data, principal component analysis (PCA) was conducted. The first 5 principal components explaining 85.8% of the variance in the data were used for PAMs, and the first 3 principal components explaining 84.9% of the variance in the data were used for PPL1s, to build the models. Each trial was indexed as an active, resting (inactive for less than 1 min) or sleep trial (inactive for more than 5 min). The state was determined from the fly’s behaviour during the period up until the onset of odor application. The same number of trials from each state were used to train the model (125 trials for each state, corresponding to 70% of all the trials in the active state, which had the least trials). Testing was done on the remaining data for each state. This procedure was repeated 50 times, using random sampling of training and test data each time, with the final accuracy of prediction for each state taken as the means.

For decoding of behavioural states from KC baseline activity, a linear discriminant decoder was used (due to lower numbers of trials/class) to discriminate the 3 states, using data from 4 flies that slept at least once during the experiment (non-sleeping flies were excluded). Baseline activity was processed as described earlier for each fly, following which data from 8 randomly selected KCs were pooled for each fly (the number 8 was selected to approximate the dimension of compartments used in PAM/PPL1 decoding analysis). PCA was conducted and the first 6 principal components explaining 86.8% of the variance in the data were used to build the model. As described for DAN analysis, behavioural states were indexed and the number of training trials was balanced across states (15 training trials from each state were used to train the model, corresponding to 60% of all the trials in the sleep state, which had the least trials). The final accuracy of prediction for each state was taken as the mean of 50 repeats of the decoding (10 repeats of random KC selection x 5 repeats of random selection of training and test data sets).

Flies were excluded from decoding analysis if their brain imaging did not result in high quality registered images with the same cell bodies or compartments traceable throughout the experiment.

#### Behavioural analyses

Behavioural analysis was performed using the output from Fictrac software ^13^, which calculated ball rotations along the X, Y, and Z axes, as well as the walking speed. Rare timepoints in which the instantaneous walking speed was unrealistically high (above 50 mm/s) were not included in the behavioural or combined brain imaging analyses, as they appeared only when Fictrac software crashed.

To define behavioural states, behaviour was manually annotated as either active or rest in Fictrac videos (each 3-5 min). Walking speeds calculated by Fictrac during the annotated periods were plotted as a histogram (Fig. 1b), allowing us to distinguish active from rest using a threshold of 1.2 mm/s (with moving average of 5 s). This threshold was subsequently used to define periods of rest and walking. Sleep was identified as periods during which the fly remained resting (inactive) for at least 5 min. To examine state-dependent differences in neural activity during odor application, the behavioural state was defined at the time of odor offset (to ensure the fly was still in that state during the odor application period), whereas for analysis of state-dependent behavioural responses, sleep and wake states were defined at the time of odor onset (to examine the behavioural response evoked by odors). In Fig. 1d, the percentage of trials in which each fly responded to the scan start of the laser was calculated. A response was scored when the fly significantly changed the walking speed (above the walking threshold of 1.2 mm/s) when comparing 4 s before and after the scan start. Similarly, in Fig. 2c and Fig. 3c, arousal to odors was analyzed in the same way, comparing 4 s before and after the odor application period.

To examine the relationship between DAN activity and spontaneous behaviour during wakefulness, the mean walking and turning speeds were correlated with mean DAN activity during 6 s of baseline period prior to odor application, only in trials where flies were awake during the analyzed period. Data were Z-scored for each fly and pooled to acquire the Pearson’s correlation and p value (with significance adjusted for multiple comparisons in the 15 DAN compartments with Bonferroni correction). Forward walking speed was calculated in trials where flies were walking forward (as opposed to backward), and forward turning was calculated as angular velocity in trials where flies were walking forward above 15°/s (to examine turning as a more goal-directed, search-like behaviour as opposed to an avoidance-like turning). Backward walking was calculated as backward velocity and backward turning was calculated as angular velocity in trials where flies were walking backward.

For quantification of behavioural responses upon arousal to odors, trials in which resting or sleeping flies significantly responded to odors (with an increase in walking speed) were examined (Fig. 4). Responses to a particular odor were included in the analysis only when they were collected in both rest and sleep states in the same fly. Due to the rarity of such events, the sample size was increased by combining data from all our data sets (four sets of experiments) in which 16 odors were used. For consistency, data corresponding to 4 s before and 4 s after the odor onset were used to assess the state and behavioral responses for each dataset (the duration of odor application was different between experiments as described earlier). To analyze turning and forward behaviour, walking speed was smoothed by a 0.5 s moving window for noise reduction.

#### Statistics

Sample sizes were predetermined based on the effect size and variability of data observed in pilot experiments. Data were assessed for normality using the Lilliefors test prior to using statistical tests (in paired statistical tests, a paired t-test was used if both data sets were normally distributed, whereas a Wilcoxon signed-rank test was used if one or both of the datasets were non-normally distributed). Corrections for multiple comparisons were performed where appropriate. Neither randomization nor blinding was performed during experiments or data analysis.

## Acknowledgements

We thank Y. Aso, A. Lin, A. Fiala, H. Tanimoto, A. Claridge-Chang, and the Bloomington Stock Center for fly stocks, T. Bockemühl for help with designing the fly ball-holder, members of the Kazama laboratory for their support and comments on the manuscript, and B. van Swinderen and J. Johansen for their comments on the manuscript. This work was supported by the Human Frontier Science Program (HFSP) fellowship to L.K., grants from Japan Society for the Promotion of Science (KAKENHI grant number JP19F19718 to L.K., JP18H02532, and JP21H04789 to H.K.), Japan Science and Technology Agency (CREST grant number JP24028095), Toray Science Foundation (23-6402), RIKEN, Kao Corporation, and Asahi Corporation to H.K.

## Author information

Contributions: L.K and H.K conceived the study and wrote the manuscript. L.K. collected and analyzed the data. M.S. assisted with setting up imaging equipment and wrote code for image registration.

## Ethics declarations

Competing interests: The authors declare no competing interests.

